# Sustained exposure to multivalent antigen-decorated nanoparticles generates broad anti-coronavirus responses

**DOI:** 10.1101/2024.10.01.616060

**Authors:** Julie Baillet, John H. Klich, Ben S. Ou, Emily L. Meany, Jerry Yan, Theodora U. J. Bruun, Ashley Utz, Carolyn K. Jons, Sebastien Lecommandoux, Eric A. Appel

## Abstract

The threat of future coronavirus pandemics requires developing cost-effective vaccine technologies that provide broad and long-lasting protection against diverse circulating and emerging strains. Here we report a multivalent liposomal hydrogel depot vaccine technology comprising the receptor binding domain (RBD) of up to four relevant SARS and MERS coronavirus strains non-covalently displayed on the surface of the liposomes within the hydrogel structure. The multivalent presentation and sustained exposure of RBD antigens improved the potency, neutralizing activity, durability, and consistency of antibody responses across homologous and heterologous coronavirus strains in a naïve murine model. When administrated in animals previously exposed to the wild-type SARS-CoV-2 antigens, liposomal hydrogels elicited durable antibody responses against the homologous SARS and MERS strains for over 6 months and elicited neutralizing activity against the immune-evasive SARS-CoV-2 variant Omicron BA.4/BA.5. Overall, the tunable antigen-decorated liposomal hydrogel platform we report here generates robust and durable humoral responses across diverse coronaviruses, supporting global efforts to effectively respond to future viral outbreaks.

**Progress and Potential:** Rapidly mutating infectious diseases such as influenza, HIV, and COVID-19 pose serious threats to human health. Yet, most vaccines still do not mount durable protection against mutagenic viruses and fail to induce broad responses to protect against emergent strains. Materials approaches to vaccine design, such as employing sustained delivery approaches or decorating nanoparticle constructs with multiple antigens, have shown promise in improving the breadth and potency of vaccines. Yet, these approaches typically require cumbersome chemistries and have not been explored in pre-exposed populations over clinically relevant time scales. Here, we report the development of an injectable liposomal hydrogel depot technology capable of prolonged presentation of multiple coronavirus antigens non-covalently coordinated on the surface of the liposomes forming the hydrogel structure. These hydrogels improve the potency, durability and breadth of vaccine response and are easy to fabricate, enabling the rapid design of next generation vaccines that confer protection against rapidly evolving pandemics.

## Introduction

Scientific innovations over the last century have propelled the development of next-generation vaccine technologies to tackle severe infectious diseases, as recently shown by the FDA approval of mRNA COVID-19 vaccines. Although effective in reducing COVID-19 infections and serious health issues, mRNA vaccines face several challenges in terms of cost-effectiveness, broad efficacy, and durability.^1^ The emergence of more infective and immune evasive variants such as Omicron BA.4/BA.5 or XBB1.5^2^ and the threat of future outbreaks arising from other highly pathogenic betacoronaviruses such as SARS-CoV-1 or MERS-CoV prompt the demand for broadly protective vaccine technologies. To date, roughly a third of COVID-19 vaccine candidates in clinical development leverage SARS-CoV-2 viral protein antigens and adjuvants on account of their proven safety, stability, and affordability.^3^ Literature reports have shown that controlled spatiotemporal presentation and delivery of antigens and adjuvants could have tremendous effects on the magnitude and quality of immune responses, leading to the surge of innovative biomaterials strategies for vaccine delivery.^4–18^ Yet, developing cost-effective and scalable delivery technologies that generate potent, durable, and diverse antibody responses providing protection against currently circulating and future variants remains a technological challenge.

In addition to designing potent adjuvant systems by encapsulation, surface presentation, sustained delivery, or functionalization to polymers,^19–24^ engineering systems providing biomimetic antigen presentation has emerged as a strategy to augment long-lasting immunity, breadth, and cross-reactivity.^25–29^ Owing to their key role in mediating viral entry and the infection of host cells, the receptor binding domain (RBD) of the Spike trimer of SARS-CoV-2 has been identified as key targets against which to mount an effective immune response. For example, the multimeric display of identical RBD units on the surface of nanoparticles has been shown to significantly improve the production of highly specific neutralizing antibodies compared to those generated by soluble or encapsulated counterparts due to enhanced presentation and lymphatic uptake.^30–34^ In an effort to address the potential for future coronavirus outbreaks, several reports have shown that the combination of distinct coronavirus viral strains was instrumental in generating a greater diversity of cross-reactive antibodies that could target conserved RBD epitopes.^35–49^ This unprecedented level of cross-reactivity led to complete protection against homologous and heterologous betacoronavirus challenges.^44,47^ Thus, multivalent display approaches show great promise as pan-coronavirus vaccines, yet several challenges remain to be addressed. Most reported antigen-nanoparticle couplings make use of chemically modified protein antigens covalently attached to nanoparticles, which can hinder scale-up and their versatility. Furthermore, the seroprevalence of the SARS-CoV-2 virus, up to 96.4% in adults aged 16 years and older in 2022 through previous infection or vaccination, raises questions about the efficacy of current approaches to generate broad responses against emerging variants of concern in a population with pre-existing immunity.^50^

Across existing therapeutic nanocarriers, liposomes have been widely used in clinical drug product formulations due to their ease of manufacture, biomimetic nature, bioactivity and versatility.^51^ Notably, commercially available phospholipids functionalized with a Cobalt NTA chelator arm can non-covalently bind with exceptional affinity (K_d_ ∼ 10^−13^ M) to His-tagged regions commonly functionalized to recombinant proteins such as subunit antigens.^52^ Hence, formulating liposomes decorated with Co-NTA ligands or derivatives offers a unique and modular lever to display diverse antigens without chemical alteration.^34,53^ While chelating liposomes afford a promising approach to enhance spatial control over antigen presentation, we and others have shown that sustaining the (co)-exposure of antigens and adjuvants timeframes is also pivotal for enhancing germinal center activity, antibody affinity, durability, and the diversification of the humoral immune response.^12,18,54–58^ In this regard, our group previously showed that Co-NTA decorated liposomes could be used as a structural motif to form supramolecular hydrogels when mixed with modified biopolymers and enable prolonged release of model proteins *in vitro* and *in vivo*.^59^ These liposomal hydrogels are injectable, biocompatible, resorbable over several weeks (tunable with hydrogel formulation), and do not evoke foreign body responses in rodents.

Here, we sought to develop a liposomal hydrogel depot technology for multivalent coronavirus subunit vaccine candidates to enhance the multivalent display and prolong the exposure to distinct coronavirus strains to enhance breadth, cross-reactivity, and durability of protection. We first characterized the hydrogels’ mechanical properties and demonstrated that liposomal hydrogels allow favorable immune cell infiltration *in vivo*. We then assessed the humoral and cellular immune responses generated by vaccination primed with liposomal hydrogels decorated with wild-type (WT) RBD and boosted with a multivalent presentation of WT and BA.4/BA.5 RBDs compared to soluble clinical controls. We demonstrated that hydrogel vaccination led to consistently higher and more durable antibody titers against homologous and heterologous antigens, with substantial improvements in complete and rapid seroconversion, over soluble controls. These results were consistent with our findings of more active germinal centers and increased memory B cell populations in mice immunized with hydrogel vaccines. We corroborated these findings with a more intricate mosaic liposome display consisting of four coronavirus strains or variants of interest, demonstrating similar improvements in breadth, durability, and potency of humoral and cellular immune response in mice vaccinated with the hydrogel vaccines compared to clinical controls. Finally, we explored a tetravalent hydrogel vaccination in a pre-exposed model where mice were first immunized with a standard soluble prime-boost vaccine comprising Hexapro, a Spike variant from WT SARS-CoV-2, before receiving a boost of tetravalent hydrogel six months later. Pre-exposed mice immunized with tetravalent hydrogel vaccines demonstrated more robust and long-lasting antibody titers against diverse strains, as well as neutralizing antibodies against WT and BA.4/BA.5, than control immunizations. Together, we demonstate that multivalent liposomal nanoparticle hydrogel (LNH) vaccines are promising candidates for overcoming challenges in designing pan-coronavirus vaccines.

## Results

### Formulation and characterization of antigen-liposomal hydrogels

To explore the outcome of multivalent antigen presentation and extended vaccine exposure on immune responses, we designed a subunit coronavirus LNH vaccine candidate consisting of diverse coronavirus strains combined with a molecular adjuvant (Figure 1a). We previously reported the self-assembly of LNHs through the dynamic non-covalent interactions between biopolymers functionalized with pendant aliphatic chains and the hydrophobic membranes of liposomes.^59^ We formulated Co-NTA decorated liposomes from a mixture of DMPC (1,2-dimyristoyl-*sn*-glycero-3-phosphocholine), DMPG (1,2-dimyristoyl-*sn*-glycero-3-phospho-(1’-rac-glycerol) sodium salt), cholesterol, and DGS-NTA(Co) (1,2-dioleoyl-sn-glycero-3-[(N-(5-amino-1-carboxypentyl)iminodiacetic acid)succinyl] cobalt salt) using the thin film rehydration – extrusion method (9:1:2:0.379 molar ratio). The resulting Co-NTA-liposomes were monodispersed with a hydrodynamic diameter averaging 102 nm by Dynamic Light Scattering (DLS; Figure 1b and S1a). Incubation with commercially available His-tagged WT SARS-CoV-2 RBD antigens allowed for non-covalent surface chelation of the RBD proteins, confirmed by an increase of 10 nm in hydrodynamic diameter. Antigen-decorated liposomes and adjuvants were then simply mixed with a solution of dodecyl-modified hydroxypropylmethylcellulose (HPMC-C_12_) to yield a supramolecular LNH (2 wt% liposomes, 4 wt% cellulose derivatives, 94 wt% buffer).

**Figure 1.**
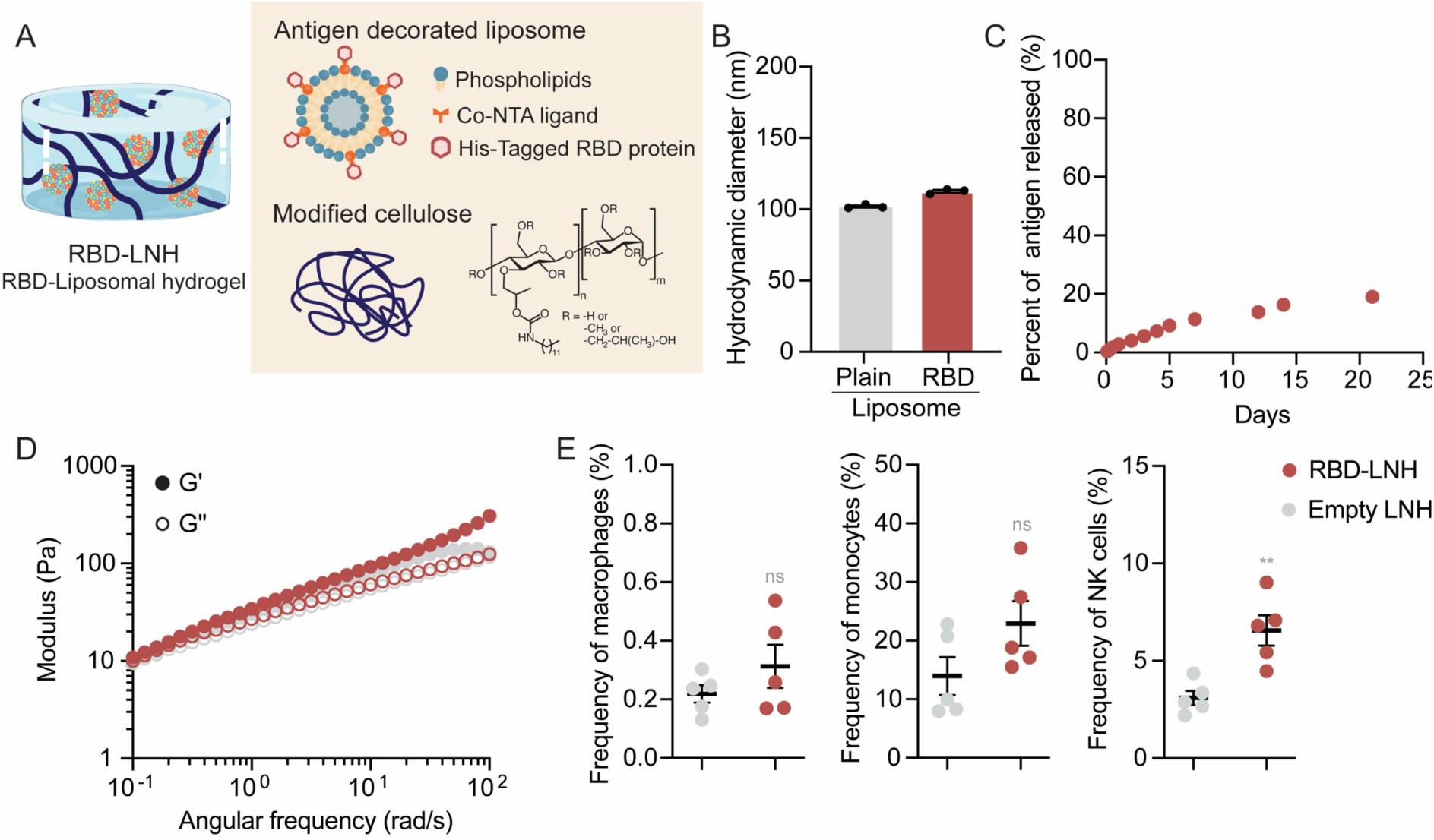
Antigen-decorated liposomal hydrogel RBD-LNH. **a.** Schematic representation of RBD-LNH consisting of His-Tagged RBD antigens chelated on the surface of liposomes interacting with cellulose derivative polymers through hydrophobic interactions. **b.** Hydrodynamic diameter of plain and RBD-decorated liposomes measured by dynamic light scattering. **c.** Percent of WT SARS-CoV-2 RBD released from RBD-LNH in vitro over 21 days. **d.** Dynamic oscillatory frequency sweep of empty and vaccine-loaded LNHs with WT SARS-CoV-2 RBD and 3M-052/AF (1% strain). **e.** Frequency of CD45^+^ leukocytes of macrophages, monocytes and NK cells infiltrating empty and vaccine-loaded LNHs (WT SARS-CoV-2 RBD, 3M-052/AF) 3 days after subcutaneous injection. See supplemental for full gating strategy. Data are shown as mean +/− SEM (n=5). *p* values were calculated from the general linear model followed by Student’s t-test and reported in Table S1.

We determined these LNH formulations had an apparent mesh size of around 4.4 nm which slows the diffusion of objects of hydrodynamic diameter of a comparable size or larger.^59^ As liposomes act as crosslinks in the hydrogel matrix, the diffusion of RBD proteins decorated on the surface of liposomes is constrained by the self-diffusion of the hydrogel matrix and are only released through erosion of the hydrogel. We investigated the kinetics of release of WT RBD antigens from LNHs with an *in vitro* capillary release model assay (Figure 1c). In these assays, RBD-LNHs were injected into capillary tubes, topped with buffer (PBS 1X), stored at physiological temperatures, and the release of cargo into the supernatant was monitored over time. The concentration of WT RBD proteins released in the supernatant over time was quantified by ELISA. After 21 days, only 20% of WT RBD proteins were released, which confirmed the immobilization and sustained exposure of antigens, dictated by the rate of erosion of LNH.

Next, we investigated whether vaccine components would alter critical physico-chemical properties of LNH such as stiffness, shear-thinning, and self-healing. We used the molecular lipidated TLR7/8a adjuvant 3M-052, which has shown to strongly enhance humoral immune responses in a previously reported COVID-19 subunit vaccine candidate.^56^ The liposomal form of 3M-052 bound to alum nanoparticles, 3M-052/AF, was selected due to its ease of immobilization into the mesh of the hydrogel. Its diffusion was thereby assumed to be slowed down to the erosion rate of LNH, similar to coordinated RBD antigens, allowing for the prolonged co-exposure of both vaccine components. Compared to empty LNH, RBD-LNH adjuvanted with 3M-052/AF exhibited overall similar rheological properties, indicating the negligible influence of the entrapped vaccine components on the LNH structure. RBD-LNH exhibited solid-like behaviors with an elastic modulus (G’) superior to loss modulus (G”) over the range of frequencies tested within the linear viscoelastic regime of the materials (Figure 1d). RBD-LNH conserved shear-thinning behaviors with increasing shear rates without rupture previously shown to enable injection through standard needles (Figure S1b), as well as self-healing properties of the LNH structure (Figure S1c). Further, these materials exhibited a yield stress of around 15 Pa, which indicates the formation of depots upon subcutaneous administration (Figure S1d).^60^ Together, these properties demonstrate that LNHs can be easily loaded with distinct cargo and can be safely injected through clinically relevant needles. Additionally, they can maintain their integrity during injection, avoiding burst release, and form depots in the subcutaneous space.

Due to the dynamics of their non-covalent crosslinks, previously described supramolecular polymer-nanoparticle hydrogel vaccines have been shown to favor the infiltration of immune cells and enhance the initiation of adaptive immune responses.^58^ Here, we quantified the type and frequency of cells infiltrating vaccine-loaded and empty LNHs three days after subcutaneous injection in mice (Figure 1e, S2 and S3, *p* values in Table S1). Overall, we found a higher frequency of antigen-presenting cells (APCs), including macrophages and monocytes, and higher or comparable frequencies of B and T cells (key mediators of adaptive immunity) in the vaccine formulations compared to empty LNH. These results suggested the incorporation of vaccine components in the LNH recruited immune cells into the hydrogel niche, driving early immune activation. We also measured a significantly higher frequency of NK cells, likely due to their activation by the potent TLR7/8a adjuvant 3M-052. NK cells have been shown to participate in the regulation of B cell and T cell responses and influence the quality of vaccine responses.^61^ Notably, their activation has been associated with robust efficacy and adaptive immunity in certain vaccines adjuvanted with lipid-based adjuvants.

### Monovalent/bivalent prime-boost RBD-LNH immunization induces potent neutralizing activity and breadth of humoral immune responses

We investigated whether the multivalent liposomal antigen presentation and prolonged exposure of vaccine components offered by RBD-LNH could improve humoral immune responses in C57BL/6 mice. The vaccination timeline consisted of a prime-boost regimen with a monovalent prime immunization of WT RBD antigen followed by a bivalent booster of WT and Omicron BA.4/BA.5 RBD antigens on week 8 (1:1 mass ratio, Figure 2a). This mixed prime-boost regimen was chosen to mimic exposure experienced by many people during the roll-out of the COVID-19 vaccines at the height of the pandemic. We compared the RBD-LNH formulations adjuvanted with 3M-052/AF to two controls composed of the soluble RBD antigens (WT during prime, WT/BA.4/BA.5 during boost) adjuvanted with clinically relevant AddaVax or 3M-052/alum adjuvants. Mice were subcutaneously injected with 100 μL of each formulation (10 μg antigen, 1 μg 3M-052 per dose) and IgG antibody titers against the homologous WT and BA.4/BA.5 antigens were monitored over 12 weeks. RBD-LNH vaccination elicited a strong and robust antibody response against the two homologous antigens tested across 12 weeks, averaging end-point titers of 10^3–5^ post-prime and 10^5–6^ post-boost against WT and BA.4/BA.5 strains (Figure 2b, *p* values in Table S2). Notably, anti-WT RBD endpoint titers generated by RBD-LNH post-prime were in the same order of magnitude as those produced from mice post-boost when vaccinated with the clinical AddaVax control.

**Figure 2.**
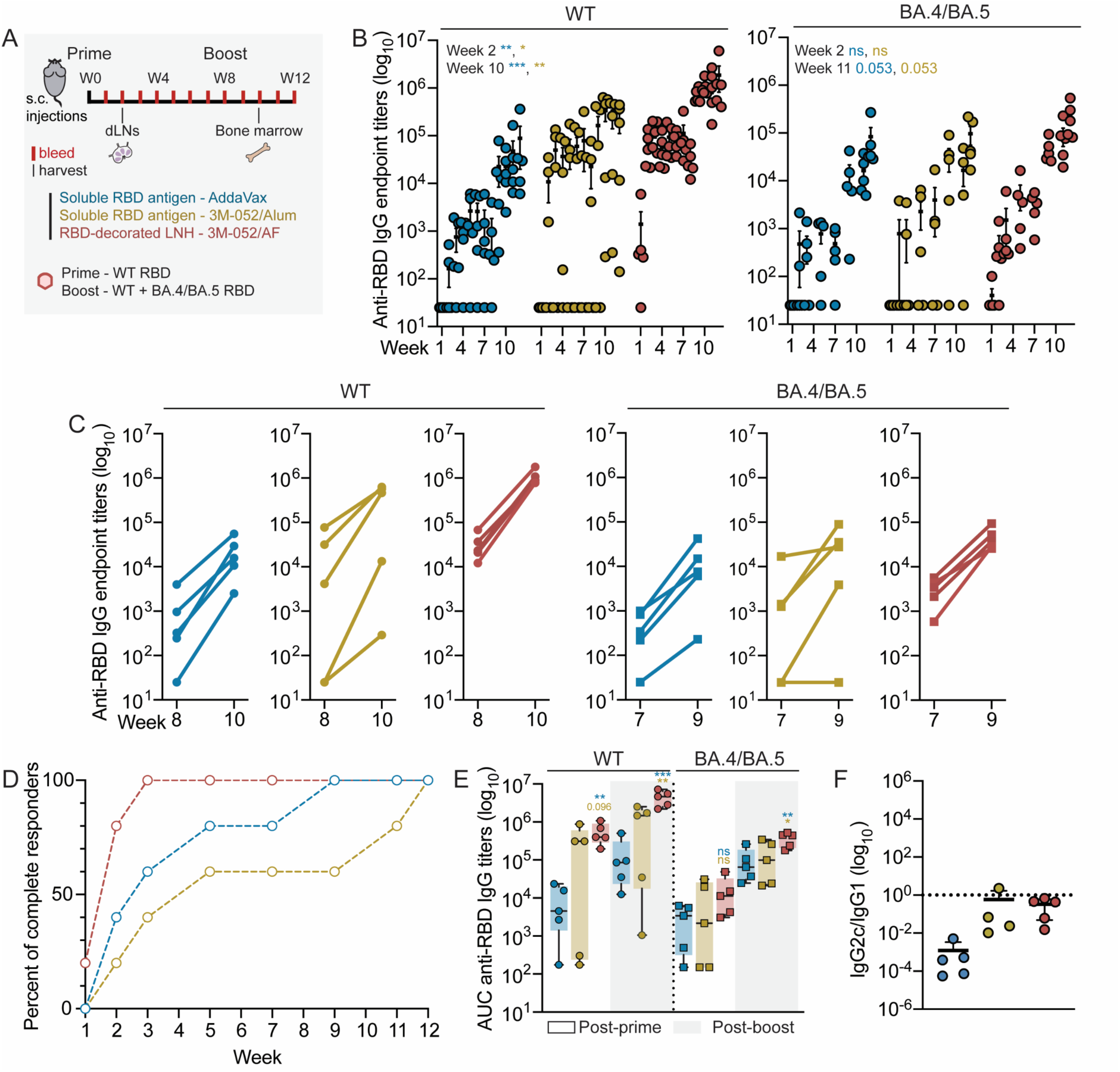
Humoral antibody response to a SARS-CoV-2 monovalent prime – bivalent boost immunization in naïve mice. **a.** Schematic of the vaccination timeline, which consisted of a monovalent WT SARS-CoV-2 RBD priming (W0) followed by a bivalent WT/BA.4/BA.5 SARS-CoV-2 RBD booster (1:1 mass ratio, W8) of RBD-LNH adjuvanted with 3M-052/AF or the soluble antigens adjuvanted with AddaVax or 3M-052/Alum controls. **b.** Anti-RBD IgG antibody endpoint titers measured over 12 weeks against WT and BA.4/BA.5 strains. **c.** Anti-RBD IgG antibody endpoint titers across mice pre and post boost at weeks 8 and 10, and 7 and 9 for WT and BA.4/BA.5 strains respectively. **d.** Percent of total mice responders showing detectable antibody endpoint titers superior or equal to 10^2^ against both WT and BA.4/BA.5 variants. An animal is considered as non-responder if quantitative titers were not measured against all the strains. **e.** Area under the curves (AUCs) of anti-RBD IgG endpoint antibody titers over 8 (post-prime) and 12 (post-boost) weeks for WT and BA.4/BA.5 strains. **f.** Ratio of anti-RBD (WT) IgG2c to IgG1 antibody endpoint titers (week 10). Data are shown as mean +/− SEM (n=4-5). AUCs are represented as box and whiskers plots showing median, interquartile range (box) and minimum/maximum values (whiskers). *p* values comparing RBD-LNH vaccination to the two soluble controls were calculated from the general linear model followed by Student’s t-test and reported in Table S2-S3.

RBD-LNH vaccines generated less overall variability in titers across mice for both antigens evaluated. Indeed, all mice immunized with RBD-LNH vaccines consistently and strongly responded to the boost performed at week 8, after which we observed that endpoint titers not only rapidly increased for both antigens but also remained clustered within an order of magnitude of each other (Figure 2c). These observations were drastically different with the two soluble controls for which the antibody titers produced spanned across four orders of magnitude pre- and post-boost, highlighting the variability induced by the prime and the lower benefit of the boost. Strikingly, RBD-LNH vaccination also induced 80% of mice to produce quantitative endpoint antibody titers equal to or above 10^2^ against both WT and BA.4/BA.5 antigens two weeks post-immunization followed by 100% of total responders by week 3 (Figure 2d). By comparison, groups vaccinated with the soluble controls adjuvanted with AddaVax and 3M-052/Alum did not produce cohorts with 100% responders until well after boosting, at weeks 9 and 12, respectively. The area under the curves (AUCs) of RBD-LNH post-prime and post-boost confirmed its superiority and robustness compared to the soluble controls (Figure 2e, *p* values in Table S3). The robust antibody response elicited by RBD-LNH against BA.4/BA.5 pre-boost was even more remarkable considering this antigen was not included in the vaccine formulations during priming, which suggests an overall enhancement in breadth conferred by the RBD-LNH vaccines.

In line with the enhanced antibody titers described above, RBD-LNH vaccines induced significantly higher IgG1 and IgG2c isotype titers at week 10, resulting in a ratio of IgG2c/IgG1 close to 1 (Figure 2f and S4, *p* values in Table S4). The soluble 3M-052/Alum control generated similar results while soluble AddaVax resulted in a ratio below 1 due to low levels of IgG2c endpoint titers. The measure of IgG isotypes and IgG2c/IgG1ratio is a common benchmark associated with Th2 and Th1 responses, suggesting a more Th1- or Th2-skewed response when the ratio is above or below 1, respectively. While soluble AddaVax controls resulted in a Th1-skewed response, 3M-052/Alum and RBD-LNH led to a balanced Th2/Th1 response which has been shown to be beneficial in COVID-19 disease outcomes.^62,63^ Notably, RBD-LNH generated less variable IgG1 and IgG2c titers resulting in a more robust and balanced Th2/Th1 response compared to 3M-052/Alum, highlighting once again the benefits of LNH in eliciting more consistent antibody responses.

We then assessed the neutralizing activity of antibodies produced from the different immunizations by measuring the infectivity of spike-pseudotyped lentiviruses into HeLa cells overexpressing the angiotensin-converting enzyme 2 (ACE2) surface receptor (Figure 3a and S5). The neutralization titer (NT_50_) was defined as the serum dilution concentration at which 50 % virus infection was prevented, hence higher NT_50_ denotes better neutralizing antibodies activity. Four weeks following RBD-LNH vaccination, we measured robust NT_50_ against WT RBD with four mice out of five producing neutralizing antibodies (*p* values in Table S5). In contrast, no neutralizing activity was observed from mice that received the soluble AddaVax control and only one responder was reported from the soluble 3M-052/Alum adjuvanted vaccine. This observation was even more striking four weeks post-boost as significantly higher neutralization titers were observed for all mice following the bivalent RBD-LNH booster compared to the two soluble controls. Moreover, four mice out of five vaccinated with RBD-LNH produced neutralizing titers to the immune evasive Omicron BA.4/BA.5 RBD variant four weeks post-boost compared to two and zero mice vaccinated with the soluble AddaVax and 3M-052/Alum controls, respectively.

**Figure 3.**
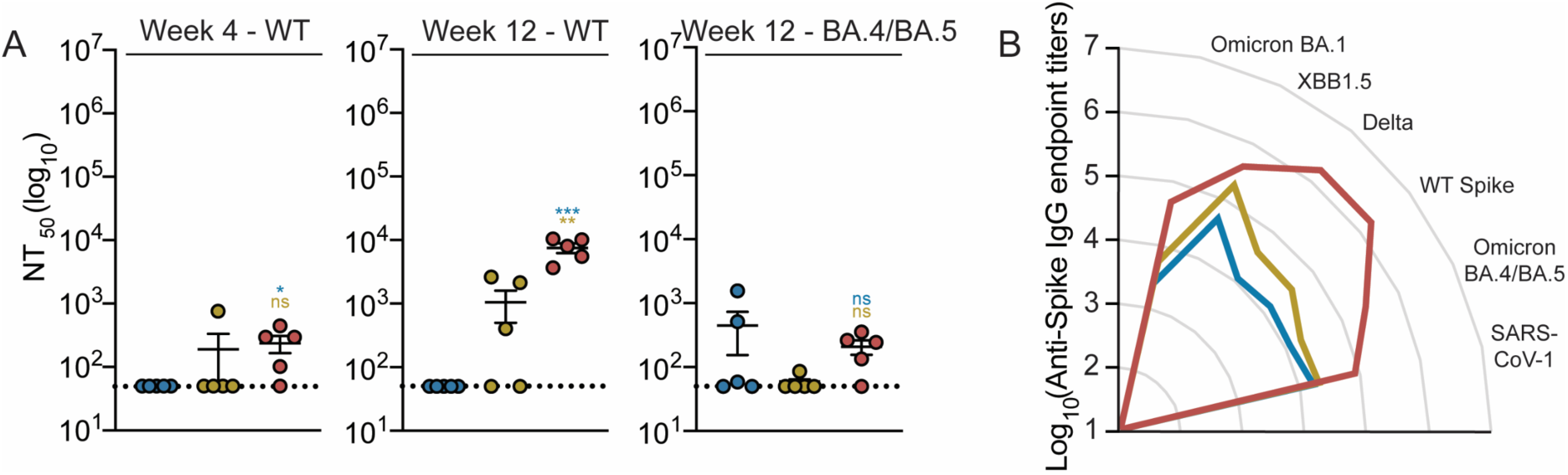
Neutralization and breadth elicited by the monovalent prime – bivalent boost immunization. **a.** Neutralizing activity of sera from immunized mice against WT and BA.4/BA.5 SARS-CoV-2 pseudoviruses, pre- and post-boost (W4, W12) represented as NT_50_ values. **b.** Anti-Spike IgG endpoint titers measured for Omicron BA.1, XBB1.5, BA.4/BA.5, Delta, WT SARS-CoV-2 and SARS-CoV-1 4 weeks post-boost (W12). Data are shown as mean +/− SEM (n=5). *p* values comparing RBD-LNH vaccination to the two soluble controls were calculated from the general linear model followed by Student’s t-test and reported in Table S5-S6.

We further explored whether RBD-LNH could enhance the breadth against heterologous variants and strains of coronaviruses of interest (Figure 3b and S6). We measured endpoint titers against sarbecoviruses of interest such as the SARS-CoV-2 variants Delta, Omicron BA.1 and XBB 1.5 as well as SARS-CoV-1 four weeks post-boost. RBD-LNH led to significantly higher titers compared to the soluble AddaVax control and overall higher responses compared to the soluble 3M-052/Alum treatment (*p* values in Table S6). RBD-LNH showed consistent antibody responses across homologous and heterologous strains and outperformed the soluble controls exhibiting high variability.

Additionally, we measured germinal center (GC) activity in the vaccine draining lymph node (dLN) by quantifying antigen-specific germinal center B cells as well as T follicular helper cells (Tfh cells) as these two populations have been shown to correlate with higher antibody affinity maturation, neutralization and memory B cell selection in COVID-19 vaccines.^6,64^ Two weeks post-prime, mice vaccinated with RBD-LNH had the highest count of germinal center B cells (GCBCs) and antigen-specific GCBCs compared to the two soluble controls (Figure 4a,b and S7, *p* values in Table S7). Moreover, mice vaccinated with RBD-LNH resulted in significantly higher count of Tfh cells (Figure 4c, *p* values in Table S7). These results indicate the superiority of RBD-LNH in generating robust GC activity and are consistent with the overall improved antibody response, neutralization, and breadth.

**Figure 4.**
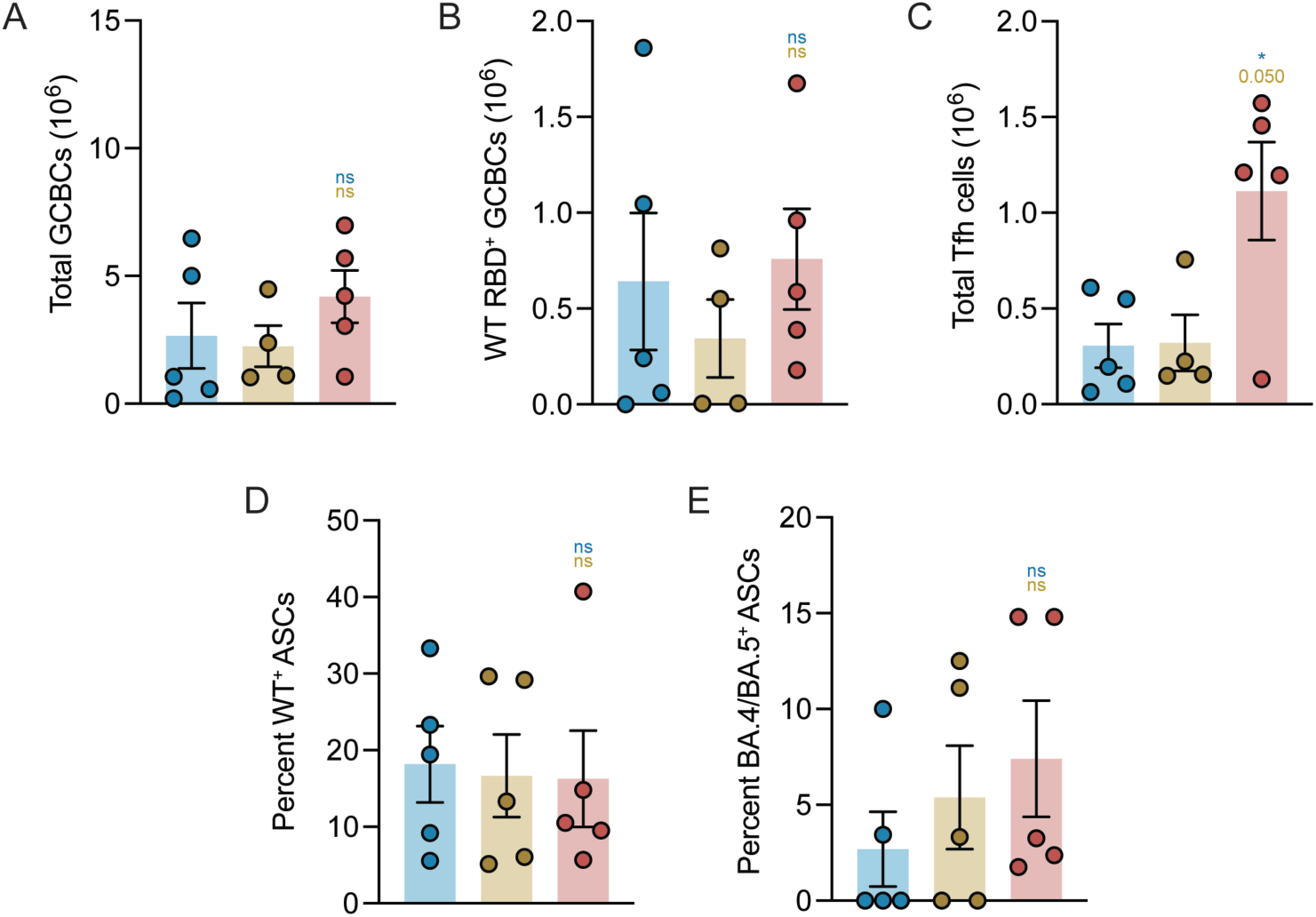
B cell responses to LNH immunization. **a.** Total germinal center B cells (GCBCs), **b.** WT RBD specific GCBCs, and **c.** total T follicular helper cells (Tfh) from the draining lymph nodes, elicited two weeks post priming immunization (W2) with WT RBD-LNH adjuvanted with 3M-052/AF or the soluble WT antigen adjuvanted with AddaVax or 3M-052/Alum controls. Percent of **d.** WT and **e.** BA.4/BA.5 antigen-specific antibody-secreting cells (ASCs) from the bone marrow, produced at week 10, two weeks post prime-boost immunization with a monovalent WT SARS-CoV-2 RBD monovalent prime and a bivalent WT/BA.4/BA.5 SARS-CoV-2 RBD boost. Data are shown as mean +/− SEM (n=5). *p* values comparing RBD-LNH vaccination to the two soluble controls were calculated from the general linear model followed by Student’s t-test and reported in Table S7-S8.

Waning immunity continues to exacerbate issues of providing population-level protection against infectious diseases, requiring frequent boosters or complex vaccination regimen.^65^ Mounting strong memory responses is therefore critical for next generation vaccines. To assess the memory response generated by RBD-LNH vaccination, we quantified the percent of antigen-specific antibody-secreting cells (ASCs) among all ASCs isolated from bone marrow, since bone marrow ASCs mostly consist of memory B cell populations such as long-lived plasma cells, via ELISpot. While the percent of WT-specific bone marrow ASCs was similar across all three treatment groups, the percent of BA.4/BA.5-specific ASCs in mice vaccinated with RBD-LNH was noticeably increased compared to those vaccinated with soluble controls (Figure 4d,e, *p* values in Table S8). Strikingly, only two and three out of five mice produced detectable BA.4/BA.5 specific IgG-secreting cells when vaccinated with soluble AddaVax and 3M-052/Alum controls, respectively, compared to all five out of five when vaccinated with RBD-LNH. These results were consistent with sustained and greater plasma antibody titers, suggesting improved serological memory and a more durable immune response from vaccination via RBD-LNH compared to soluble controls.

### Tetravalent RBD-LNH immunization induces consistent humoral immune responses against diverse betacoronaviruses

Given the ability of LNH formulations to induce a vaccine response characterized by robust neutralizing titers, breadth, and durability in a monovalent prime – bivalent boost SARS-CoV-2 regime, we sought to expand the LNH strategy towards a broader diversity of viruses, working towards pan-coronavirus applications. We designed a multivalent LNH formulation consisting of liposomes decorated with four betacoronavirus RBD antigens, including WT and BA.4/BA.5 SARS-CoV-2, SARS-CoV-1, and MERS-CoV (1:1:1:1 mass ratio). Following the same methodology as previously described, we formulated tetravalent RBD-LNH comprising tetravalent-decorated liposomes and 3M-052/AF adjuvant (2.5 μg of each antigen, 1 μg of 3M-052 per dose). The vaccination schedule consisted of a clinically relevant prime-boost immunization schedule of tetravalent RBD-LNH (weeks 0 and 8, Figure 5a). Tetravalent RBD-LNH was compared to two soluble controls comprising the same mixture of RBD antigens adjuvanted with AddaVax or 3M-052/Alum.

**Figure 5.**
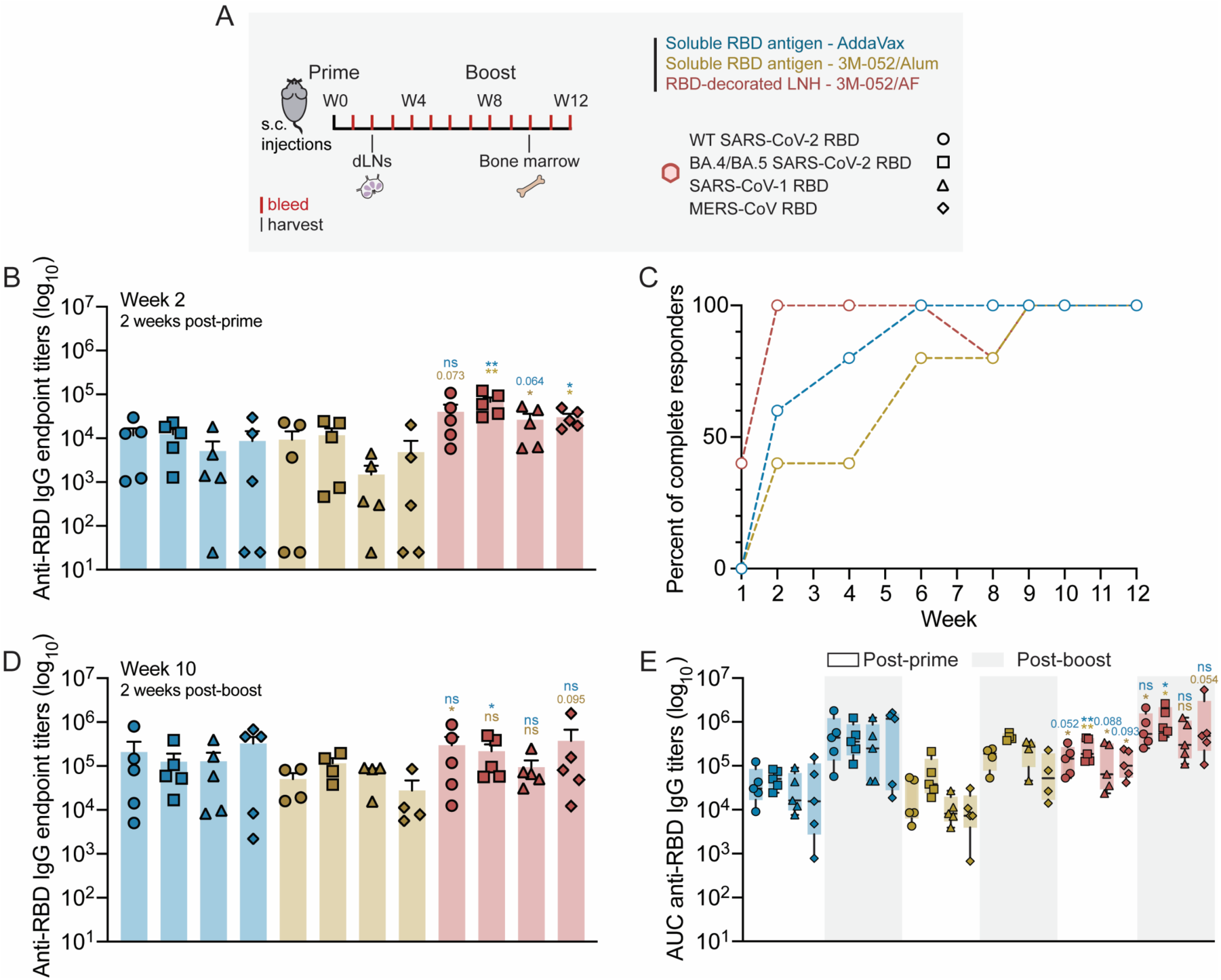
Humoral antibody response to a tetravalent betacoronavirus prime-boost immunization in naïve mice. **a.** Schematic of the vaccination timeline, which consisted of a tetravalent betacoronavirus prime-boost regime WT, BA.4/BA.5 SARS-CoV-2, SARS-CoV-1 and MERS-CoV RBDs (1:1:1:1 mass ratio; W0, W8) of RBD-LNH adjuvanted with 3M-052/AF or the soluble antigens adjuvanted with AddaVax or 3M-052/Alum controls. **b.** Anti-RBD IgG antibody endpoint titers measured two weeks post-prime (W2) against the four homologous strains. **c.** Percent of total mice responders showing detectable antibody endpoint titers superior or equal to 10^2^ against all four strains. An animal is considered as non-responder if quantitative titers were not measured against all the strains. **d.** Anti-RBD IgG antibody endpoint titers measured two weeks post-boost (W10) against the four homologous strains. **e.** Area under the curves (AUCs) of anti-RBD IgG endpoint antibody titers over 8 (post-prime) and 12 (post-boost) weeks for WT, BA.4/BA.5 SARS-CoV-2, SARS-CoV-1 and MERS-CoV strains. Data are shown as mean +/− SEM (n=4-5). AUCs are represented as box and whiskers plots showing median, interquartile range (box) and minimum/maximum values (whiskers). *p* values comparing RBD-LNH vaccination to the two soluble controls were calculated from the general linear model followed by Student’s t-test and reported in Table S9-S10.

In line with the data observed with the monovalent-bivalent RBD-LNH vaccine regime, tetravalent RBD-LNH formulations induced consistent, robust, and high antibody endpoint titers averaging 10^5^ as early as two weeks post-prime (Figure 5b, *p* values in Table S9). The two soluble control vaccinations generated a variable number of non-responders across the four strains, leading to inconsistencies. Astonishingly, the tetravalent RBD-LNH vaccine induced 100% of mice to generate quantitative titers exceeding 10^2^ against all four homologous strains by week 2 post-immunization, while the soluble AddaVax and 3M-052/Alum immunizations reached this threshold significantly later at weeks 6 and 9, respectively (Figure 5c). Two weeks post-boost, RBD-LNH immunization maintained its ability to generate overall more consistent and higher antibody titers between animals (Figure 5d, *p* values in Table S9). The AUC of antibody titers generated by mice vaccinated with tetravalent RBD-LNH further confirmed the robustness of the antibody response to all homologous strains pre- and post-boost. These data also highlight the consistency of responses across distinct viral strains such as WT and MERS-CoV compared to the soluble controls (Figure 5e and S8, *p* values in Table S10 and S11). Notably, the AUCs of tetravalent RBD-LNH vaccination titers pre-boost are comparable to those of the soluble 3M-052/Alum control post-boost for all homologous strains.

Additionally, mice immunized with tetravalent RBD-LNH vaccines showed improved quality of B cell responses. Indeed, these immunizations produced significantly higher counts of Tfh cells, total GCBCs, and antigen-specific GCBCs compared to the soluble controls two weeks post-prime (Figure 6a-c, *p* values in Table S12). Notably, although mice treated with the soluble AddaVax and 3M-052/Alum controls could produce some levels GCBCs, almost none were antigen-specific GCBCs, suggesting the ineffectiveness of the soluble vaccinations to generate robust antigen-specific B cell responses.

**Figure 6.**
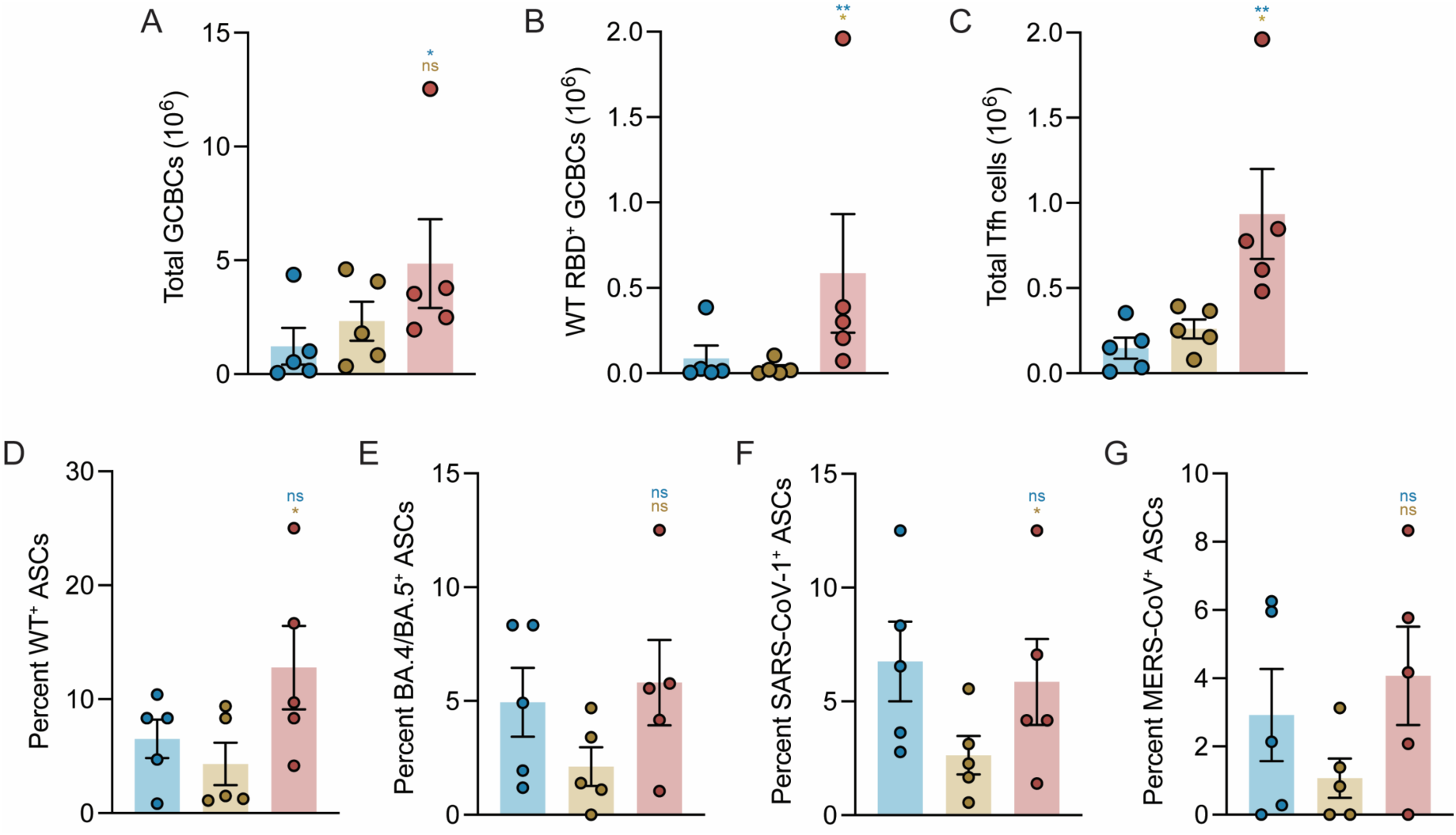
B cell responses to tetravalent LNH immunization. **a.** Total germinal center B cells (GCBCs), **b.** WT RBD specific GCBCs and **c**. total T follicular helper cells (Tfh) from the draining lymph nodes, elicited two weeks post priming immunization (W2) with tetravalent RBD-LNH adjuvanted with 3M-052/AF or the soluble antigens adjuvanted with AddaVax or 3M-052/Alum controls. Percent of **d.** WT, **e.** BA.4/BA.5, **f.** SARS-CoV-1 and **g.** MERS-CoV antigen-specific antibody-secreting cells (ASCs) from the bone marrow, produced two weeks post-boost (W10) with a tetravalent vaccine formulation. Data are shown as mean +/− SEM (n=5). *p* values comparing RBD-LNH vaccination to the two soluble controls were calculated from the general linear model followed by Student’s t-test and reported in Table S12-S13.

To further probe the breadth of memory responses induced by vaccination with tetravalent RBD-LNH, we quantified the percent of antigen-specific ASCs among all ASCs isolated from bone marrow for all four homologous antigens two weeks post-boost. We observed comparable percents of antigen-specific ASCs in mice vaccinated with soluble AddaVax control compared to tetravalent RBD-LNH and significantly greater percents of WT-specific and SARS-CoV-1-specific ASCs in mice vaccinated with tetravalent RBD-LNH compared to 3M-052/Alum (Figure 6d-g, *p* values in Table S13). Importantly, all five out of five mice immunized with tetravalent RBD-LNH vaccines had antigen-specific bone marrow ASCs against WT, BA.4/BA.5, and SARS-CoV-1 and four out of five mice had detectable specific ASCs against MERS-CoV compared to various non-responders across antigens among the soluble vaccination groups. Together, these data indicate an increased memory response in mice vaccinated with tetravalent RBD-LNH, consistent with higher circulating antibody titers, supporting the superiority of the immune response from tetravalent RBD-LNH compared to soluble controls.

### Tetravalent RBD-LNH immunization induces consistent humoral immune responses in pre-exposed animals

The prevalence of infections and exposure to the ancestral WT SARS-CoV-2 strain and variants across the global population motivates the need to pivot from the use of naïve animal models to pre-exposed cohorts. We investigated whether tetravalent RBD-LNH vaccines could generate a robust and consistent antibody response against all homologous betacoronavirus strains in mice having pre-existing immunity to the ancestral WT SARS-CoV-2 strain. To best recapitulate a lifelike vaccination schedule, we immunized naïve mice with a prime-boost regime at weeks 0 and 3 with a clinically relevant soluble vaccine formulation consisting of Hexapro Spike antigen and the TLR9 agonist adjuvant CpG adsorbed on Alum (Figure 7a). Hexapro is a stable variant of the prefusion SARS-CoV-2 spike glycoprotein, currently in phase I clinical trials by Vaxxas,^66^ and CpG/Alum is used in clinical vaccines such as in Dynavax’s hepatitis B vaccine. Six months following the initial Hexapro prime-boost regimen, we ensured mice showed comparable titers against WT SARS-CoV-2 strain (Figure S9a) and then immunized animals with either a tetravalent RBD-LNH vaccine, or one of the two soluble AddaVax and 3M-052/Alum controls. Antibody titers against the four betacoronavirus strains were measured for up to six months.

**Figure 7.**
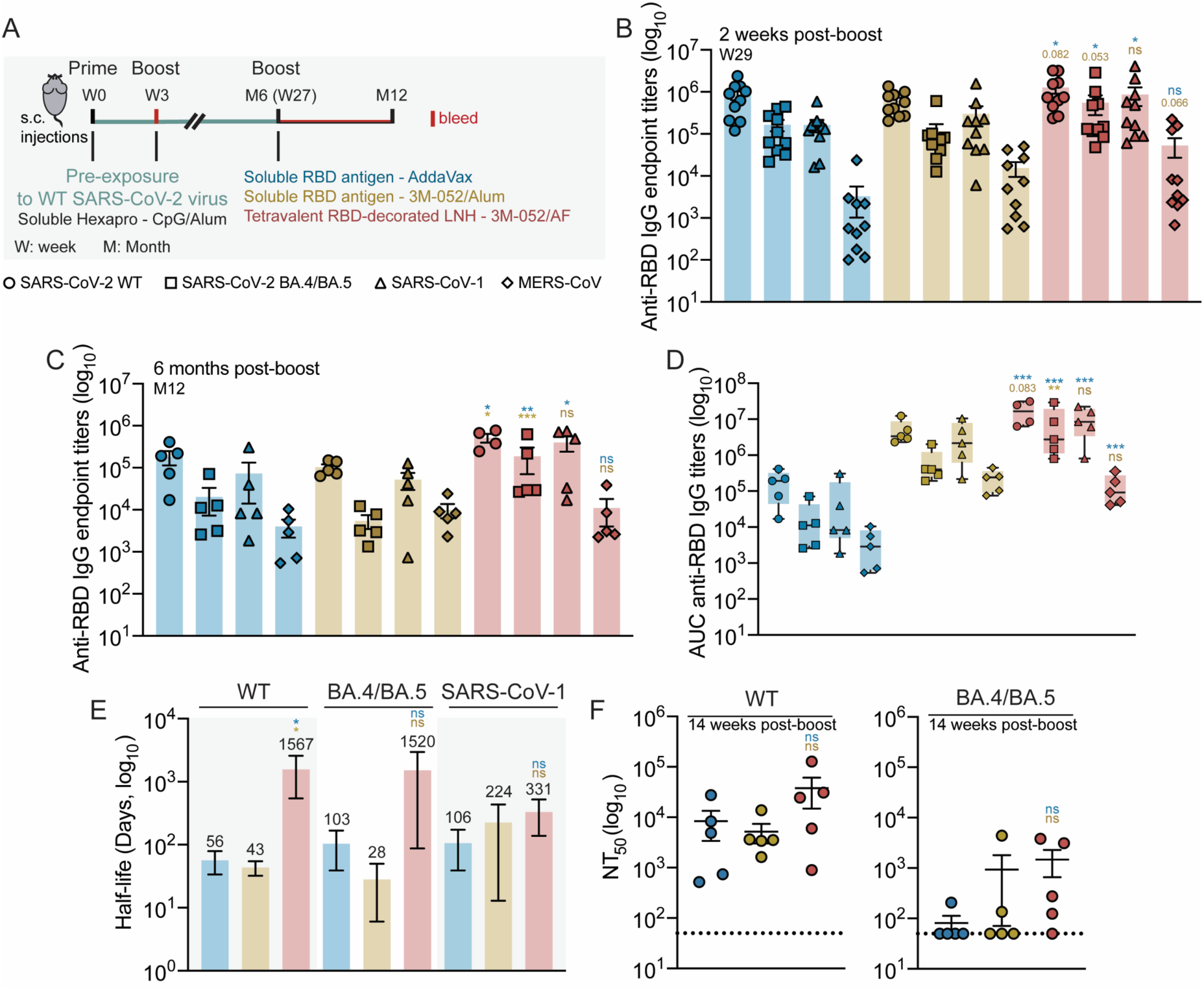
Humoral antibody response to a tetravalent betacoronavirus prime-boost immunization in mice pre-exposed to WT SARS-CoV-2 virus. **a.** Schematic of the vaccination timeline, which consisted of a prime-boost immunization of Hexapro adjuvanted with CpG/Alum (W0, W3) to generate a pre-exposure to the WT strain of the SARS-CoV-2 virus. The same cohort was then boosted with a tetravalent betacoronavirus vaccine at W27 including WT, BA.4/BA.5 SARS-CoV-2, SARS-CoV-1 and MERS-CoV RBDs (1:1:1:1 mass ratio) of RBD-LNH adjuvanted with 3M-052/AF or the soluble antigens adjuvanted with AddaVax or 3M-052/Alum controls. **b.** Anti-RBD IgG antibody endpoint titers measured two weeks post-boost (W29) against the four homologous strains. **c.** Anti-RBD IgG antibody endpoint titers measured six months post-boost (M12) against the four homologous strains. **d.** Area under the curves (AUCs) of anti-RBD IgG endpoint antibody titers over six months post tetravalent betacoronavirus vaccination for WT, BA.4/BA.5 SARS-CoV-2, SARS-CoV-1 and MERS-CoV strains. **e.** Half-life of anti-SARS antibody titers over the course of six months post tetravalent boost. **f.** Neutralizing activity of sera from immunized mice against WT and BA.4/BA.5 SARS-CoV-2 pseudoviruses 14 weeks post tetravalent boost, represented as NT_50_ values. Data are shown as mean +/− SEM (n=4-5). AUCs are represented as box and whiskers plots showing median, interquartile range (box) and minimum/maximum values (whiskers). *p* values comparing RBD-LNH vaccination to the two soluble controls were calculated from the general linear model followed by Student’s t-test and reported in Table S14-S15, S17-S18.

When boosted with the tetravalent RBD-LNH vaccine, mice showed higher antibody responses compared to the two soluble controls and comparable endpoint titers averaging 10^6^ against all SARS strains tested two weeks post-boost (Figure 7b, *p* values in Table S14). These data were significantly different from the antibody response obtained after vaccination with the soluble AddaVax and 3M-052/Alum controls, generating anti-WT RBD titers almost an order of magnitude higher than anti-BA.4/BA.5 and anti-SARS-CoV-1 responses. The superiority and consistency of the antibody response elicited by tetravalent RBD-LNH was further confirmed six months post-immunization, for which mice produced significantly higher antibody responses overall and maintained endpoint titers above 10^5^ against all SARS strains (Figure 7c, *p* values in Table S14). In contrast, antibody titers produced by mice vaccinated with either of the two soluble control formulations inconsistently dropped by over an order of magnitude in that time frame. The AUCs of the titers measured over the six-month timeframe confirmed the superiority, consistency, and durability of the titers elicited by tetravalent RBD-LNH across the distinct WT, BA.4/BA.5 and SARS-CoV-1 strains compared the soluble controls (Figure 7d and S9b, *p* values in Table S15 and S16). While titers generated against MERS were less potent overall than the SARS-related strains, tetravalent RBD-LNH booster generated endpoint titers averaging 10^5^ two weeks post-immunization, which is significantly higher than the titers elicited by vaccination with the two soluble controls. Notably, the AUC of anti-MERS titers generated from tetravalent RBD-LNH booster over six months is comparable to the AUCs of anti-WT SARS-CoV-2 produced from the clinically relevant AddaVax control.

We also assessed the half-life of antibody responses from the decrease of titers measured over the six-month period. RBD-LNH vaccines increased the half-life of anti-SARS antibody titers compared to the soluble treatments, indicating greater durability of antibody responses (Figure 7e). Specifically, LNH greatly extended the half-life of anti-WT and anti-BA.4/BA.5 SARS-CoV-2 antibody responses by a 15 to 20-fold increase while maintaining similar anti-SARS-CoV-1 antibody responses (*p* values in Table S17). We did not calculate the half-life of anti-MERS responses as the titers for some groups were close to the detection limit, rendering the measure of half-life unreliable. In addition to enhanced antibody half-life, RBD-LNH vaccines led to higher neutralization activity against both WT and the immune evasive BA.4/BA.5 Omicron variant 14 weeks post-tetravalent boost, for which four mice out of five produced neutralizing antibodies (Figure 7f and S10, *p* values in Table S18). On the contrary, while mice boosted with the soluble treatments produced neutralizing antibodies against WT, they elicited little to no neutralization activity to BA.4/BA.5.

## Discussion

The SARS-CoV-2 pandemic has triggered tremendeous advancements in modern vaccine design in record time. Nonetheless, manufacturing and storage costs of the commercial mRNA vaccines combined with the lack of durability and breadth they elicit raise global concern for future outbreaks. Immense challenges remain in vaccine equity, recognizing that one-third of the world’s population has still not received a single dose of COVID-19 vaccine.^67^ First and foremost, although the SARS-CoV-2 virus and related variants have been the central focus, the threat of pandemics arising from other highly pathogenic human coronaviruses derived from SARS-CoV-1 or MERS-CoV is gaining growing interest.^68,69^ In this work, we leveraged liposomal hydrogels enabling multivalent display and prolonged exposure of distinct betacoronavirus RBD proteins to enhance durability, breadth, neutralization, and cross-reactivity against a broad array of coronaviruses – a pivotal development toward designing pan-coronavirus vaccines that can protect against future, emerging strains.

Innovative approaches reporting the multivalent display of RBD antigens through their fusion on protein nanoparticles have shown to greatly improve cross-immunity.^40–44,46–48^ Nonetheless, the required protein design and potential generation of antibodies directed against the nanocarrier may limit broad application. We used ready-to-use LNH formulations from accessible and biocompatible compounds, commercially available His-tagged RBD protein antigens, and a molecular adjuvant, together conferring design simplicity, loading efficiency, and versatility. Notably, the non-covalent immobilization of His-tagged RBD proteins on liposomes allows for simple mixing of diverse viral strains without further design. LNH can thus uniquely be tuned and broadly applied to enhance the presentation and prolong the exposure of distinct antigens, a platform approach critically needed to face potential emerging coronaviruses outbreaks more effectively.

We showed that vaccination with multivalent RBD-LNH consistently led to more potent, durable, and cross-reactive antibody responses in naïve mice following two different immunization strategies with increased antigen diversity. In a monovalent (WT) - bivalent (WT/BA.4/BA.5) prime-boost schedule, mice vaccinated with LNH produced robust antibody titers with neutralizing activity and breadth. Notably, we showed that all mice immunized with the monovalent RBD-LNH prime elicited quantitative levels of antibodies against the homologous WT and heterologous BA.4/BA.5 strains as early as three weeks post-immunization. Moreover, all showed high neutralization levels to WT and four out of five elicited neutralizing antibodies to BA.4/BA.5 post-boost. These observations were remarkable considering the immune-evasive nature of the BA.4/BA.5 Omicron variant, which is known to lower the levels and efficacy of anti-BA.4/BA.5 antibodies. The improved breadth conferred by LNH against BA.4/BA.5 was further confirmed against relevant sarbecovirus strains of interest such as BA.1, Delta, XBB1.5 and SARS-CoV-1. In contrast, soluble vaccines adjuvanted with clinically relevant AddaVax or 3M-052/Alum lacked overall efficacy as shown by high disparities where 20 and 40%, respectively, of mice did not respond pre-boost and low neutralizing activity post-prime and boost. When used in a more intricate tetravalent display of the betacoronavirus WT, BA.4/BA.5, SARS-CoV-1, and MERS-CoV strains, LNH elicited identical strong features. Notably, we showed rapid seroconversion against all four homologous strains two weeks post priming immunization as well as robust and comparable antibody responses across animals and viral strains. In contrast, soluble controls showed variability as total seroconversion was only observed significantly later, close to the boost or post-boost resulting in titers spanning up to two orders of magnitude across animals vaccinated with the same treatment.

To further evaluate the breadth and potency of the immune response driven by LNH vaccination, we examined the germinal center activity in the draining lymph nodes. Since prolonged and robust GC activity is indicative of greater affinity maturation, potentially modulating more potent responses against less immunodominant antigen epitopes, we quantified the number of germinal center B cells, RBD-specific GCBCs, and number of T follicular helper cells in the draining lymph nodes. Across these metrics, we showed that germinal centers in mice vaccinated with LNH trended larger by GCBC count, contained more antigen-specific GCBCs, and contained statistically more T follicular helper cells, essential for B cell help, than those in mice vaccinated with soluble controls.

Additionally, among growing concerns about emerging variants and durability of protection, the ability of a vaccination platform to generate not only transient humoral immunity, as measured by plasma antibody titers, but also a long-term memory response is of utmost importance. We determined the percent antigen-specific antibody-secreting cells among bone marrow memory cell populations, and we showed that despite the presence of similar percents of WT RBD-specific ASCs, mice vaccinated with LNH generated more BA.4/BA.5 ASCs compared to those vaccinated with soluble controls. Impressively, all five mice generated detectable BA.4/BA.5 specific ASCs when vaccinated with LNH whereas only 40% and 60% of mice from the soluble AddaVax and 3M-052 controls generated detectable BA.4/BA.5 specific ASCs.

We demonstrated great benefits associated with LNH vaccination in naïve mice to generate potent, consistent, durable, and broad antibody responses against betacoronaviruses. Yet, a large fraction of the world population has already been exposed to the SARS-CoV-2 virus through vaccination or infection, leading to pre-existing immunity to the ancestral WT strain or related variants. Very few reports have attempted to confirm whether broad antibody responses observed in naïve murine models are seen in pre-exposed animal cohorts.^48^ Bjorkman, Townsend, Howarth and collaborators recently showed that RBD quarter nanocages could elicit broad antibody responses against diverse homologous and heterologous sarbecovirus strains three weeks following a priming immunization with Hexapro WT spike protein. Yet, important features regarding mixture of more genetically diverse viral strains and repetitive SARS-CoV-2 exposure over clinically relevant timescales remain to be explored. While SARS-CoV-2 shares 79% genetic similarities with SARS-CoV-1, it drops to 50% when compared to MERS-CoV.^70^ Martinez, Baric, Haynes, Saunders and collaborators showed that the co-presentation of all three betacoronaviruses (SARS-CoV-2, SARS-CoV-1, MERS-CoV) conferred protection following sarbecovirus and merbecovirus challenges in naïve mice while a monovalent SARS-CoV-2 display was effective against sarbecovirus challenge only.^44^ We immunized mice that had previously received two vaccinations of Hexapro, similar to the clinical primary series of mRNA vaccinations, with the tetravalent betacoronavirus RBD-LNH vaccine six months later. Antibody responses against all four homologous strains measured over six months revealed remarkably strong, consistent, and durable antibody responses, specifically against SARS-related strains. Both soluble controls failed to maintain broad antibody responses as vaccinated mice showed higher anti-WT antibody titers, seemingly narrowing the antibody response to the WT strain. Moreover, tetravalent RBD-LNH vaccination showed robust durability with a great increase of anti-SARS antibody half-life, while titers elicited from soluble vaccinations waned rapidly. The antibody response against MERS-CoV following RBD-LNH immunization was lower than the one observed for SARS-related strains possibly due to higher antigenic and genetic diversity. Nonetheless, the AUC of anti-MERS-CoV response was comparable to the one of anti-WT titers generated by the clinically relevant soluble AddaVax formulation, revealing enhanced persistence of antibody response from genetically diverse coronavirus strains following RBD-LNH vaccination. Additionally, RBD-LNH enhanced the production of neutralizing antibodies not only to WT but also BA.4/BA.5, which has been reported to be lowered in human and murine cohorts with prior infection or vaccine-induced immunity to the ancestral strain of the SARS-CoV-2 virus.^71–73^

Overall, we demonstrated that LNH is a highly modular platform that can be easily loaded with a range of genetically diverse coronaviruses, bolstering cross-reactive antibody responses and long-lasting immunity. The prolonged exposure and enhanced presentation of antigens resulted in rapid seroconversion, greater durability, and increased breadth against several distinct variants and strains, which in turn yielded better neutralization and cross-reactive responses in naïve mice. When used in murine models with pre-existing immunity to the ancestral SARS-CoV-2 strain, this technology greatly improved consistency and durability of humoral responses. Additional characterization of the repartition and density of antigens on the surface of liposomes could be explored to further optimize this platform. Moreover, literature has shown that combining too many genetically diverse RBD antigens could dilute antigen immunogenicity.^44^ Modulating the ratio and number of antigens presented could be further investigated to improve understanding of resulting antibody responses in naive and pre-exposed cohorts.

## Conclusion

Originally found in camel and bat species, SARS-CoV-2, SARS-CoV-1, and MERS-CoV are zoonotic viruses for which spillovers have been recognized as the prevalent cause of epidemics and pandemics. In the recent years, increased contact between humans and animals has favored cross-species transmission and bolsters the risk of future viral outbreaks. Despite rapid development of effective vaccines, global access as well as broad and durable protection in a pre-immune population are still highly challenged. The multivalent liposomal hydrogel depot technology we report here is an easy-to-use and modular platform able to generate robust and durable humoral immune responses across coronaviruses in both naïve and pre-exposed animals, emphasizing its potential as a pan-coronavirus strategy to improve our response to future outbreaks. Importantly, the modularity of this approach could be extended to other viruses arising from zoonotic spillovers such as flu and even combine coronaviruses and flu viruses into a single formulation to generate broad and durable immunity against recurring and challenging viruses.

## Experimental procedures

### Resource availability

#### Lead contact

Further information and requests for resources and materials should be directed to and will be fulfilled by the lead contact, Eric A. Appel.

#### Data and code availability

Data generated to evaluate the conclusions of this article are provided within the article and supplemental information. Raw data will be made available from the lead contact upon request. This paper does not report original code.

#### Materials availability

This study did not generate new, unique reagents.

### Materials

1,2-dimyristoyl-*sn*-glycero-3-phosphocholine (DMPC), 1,2-dimyristoyl-*sn*-glycero-3-phospho-(1’-rac-glycerol), sodium salt (DMPG), Cholesterol and 1,2-dioleoyl-*sn*-glycero-3-[(N-(5-amino-1-carboxypentyl)iminodiacetic acid)succinyl], cobalt salt (DGS-NTA(Co)) were purchased from Avanti Polar Lipids. NanoSizer MINI Liposome Extruders (200 nm, 100 nm) were acquired from T&T Scientific. (Hydroxypropyl)methyl cellulose, *N*-methylpyrrolidone (NMP), *N,N*-Diisopropylethylamine (Hunig’s base), dodecyl isocyanate, and bovine serum albumin (BSA) were purchased from Sigma-Aldrich. Dialysis tubing was purchased from Spectrum Labs. Sterile PBS 1X was purchased from Thermo Fisher Scientific. His-tagged protein antigens were purchased from Sino Biological: SARS-CoV-2 Spike RBD-His Recombinant Protein (cat. Number 40592-V08H-1), SARS-CoV-2 BA.4/BA.5/BA.5.2 Spike RBD-His Recombinant Protein (cat. Number 40592-V08H130), SARS-CoV-2 Delta B.1.617.2 Spike S1 Protein (cat. Number 40591-V08H23), SARS-CoV-2 Omicron B.1.1.529 Spike S1 Protein (cat. Number 40591-V08H41), SARS-CoV-2 XBB.1.5 Omicron Spike RBD Protein (cat. Number 40592-V08H146), SARS-CoV Spike/RBD-His Recombinant Protein (cat. Number 40150-V08B2-100), MERS-CoV Spike/RBD-His Recombinant Protein (aa 367-606 fragment, cat. Number 40071-V08B1-100). TLR7/8 agonist 3M-052-aqueous formulation (3M-052/AF) and 3M-052/Alum were purchased from 3M and the Access to Advanced Health Institute. Soluble spike protein Hexapro was provided by Prof. Neil King at the Institute for Protein Design, University of Washington. CpG ODN 1826 (Vac-1826), Alum and Alhydrogel 2% (vac-alu) were purchased from Invivogen. Goat anti-mouse IgG Fc secondary antibody (A16084) HRP (Horseradish peroxidase) was purchased from Invitrogen. Goat anti-mouse IgG1 and IgG2c Fc secondary antibodies with HRP (ab97250, ab97255) and 3,3’,5,5’-Tetramethylbenzidine (TMB) ELISA Substrate, high sensitivity was acquired from Abcam. SARS-CoV-2 Spike RBD (Wild Type) Specific ELISA Kit was purchased from AcroBiosystems (cat. Number RAS-A116). Reagents used for flow cytometry, neutralization and ELISpot assays are described in the corresponding sections. Unless otherwise stated, all chemicals were used as received without further purification.

### Liposomal hydrogel formulation

Liposomal hydrogels were formulated by mixing a solution of dodecyl-modified (hydroxypropyl)methyl cellulose with a solution of liposomes and were prepared according to the literature.^59^ Briefly, (hydroxypropyl)methyl cellulose (1 g) was dissolved in anhydrous NMP (45 mL) overnight at room temperature. The solution was then heated at 50 °C for 45 mins. Once the mixture reaches the stated temperature, a solution of dodecyl isocyanate (110 mg, 0.52 mmol) in 5 mL of anhydrous NMP and Hunig’s base (10 drops) were added dropwise. After 45 mins of heating, the reaction was stirred at room temperature for 20 h. The polymer was precipitated using acetone (600 mL), redissolved in Milli-Q water (∼ 2 wt%) and dialyzed (3 kDa MWCO) against water for 4 days. The polymer solution was then lyophilized and reconstituted to a 60 mg/mL or 6 wt% solution in PBS 1X and stored at 4 C.

Liposomes were prepared via the thin-film and rehydration method with a 9:1:2:0.379 molar ratio of DMPC:DMPG:Cholesterol:DGS-NTA(Co). DMPC (76.61 mg, 3064 μL, 25 mg/ml in chloroform), DMPG (8.65 mg, 173 μl, 50 mg/ml in chloroform), Cholesterol (9.71 mg, 194 μL, 50 mg/ml in chloroform) and DGS-NTA(Co) (5.03 mg, 503 μL, 10 mg/ml in chloroform) were mixed in a 10 mL round bottom flask and solvents were slowly evaporated under pressure while stirring at 40 °C until formation of a homogeneous thin film. The latter was then rehydrated under constant stirring in 1 mL of PBS 1X for 45 mins. The solution was successively extruded through 200 nm and 100 nm filters (15 passes each). Size and polydispersity were evaluated by dynamic light scattering using a DynaPro II plate reader (Wyatt Technology).

Liposomal hydrogels were prepared by mixing appropriate amounts of modified cellulose (6 wt%) and liposomes (10 wt%) solutions in PBS 1X via two syringes connected through an elbow mixer to give a 2 wt%-4 wt% polymer-liposomes content (3:2.5 weight ratio).

### Rheological characterization

Measurements were performed on a Discovery HR-2 torque-controlled rheometer with Peltier stage (TA Instruments) at 25 °C using a 20 mm serrated plate geometry and a 500 µm gap height. Dynamic oscillatory frequency sweep experiments were conducted with a constant 1% strain and angular frequencies from 0.1 to 100 rad/s within the linear viscoelastic regime. Flow sweeps were performed with shear rates from 0.01 to 10 s^−1^. Step-shear experiments were done by altering between low shear rates (0.1 rad/s for 60 s) and high shear rates (10 rad/s for 60 s). Stress-controlled flow sweep measurements were conducted at shear rates from 0.001 to 10 s^−1^. Stress controlled yield stress measurements (stress sweeps) were performed from low to high stress with steady-state sensing and 10 points per decade.

### In vitro release assay

100 μL of a liposomal hydrogel vaccine dose loaded with WT SARS-CoV-2 RBD (10 μg) were injected at the bottom of a capillary tube plugged at one end with epoxy. The tube was covered with 400 uL of PBS 1X and incubated at 37 °C for 21 days. The supernatant was removed at each timepoint and replaced with fresh buffer. The amount of WT RBD antigen released was quantified using a SARS-CoV-2 Spike RBD (Wild Type) Specific ELISA Kit (AcroBiosystems) according to the manufacturer’s instructions. Experiments were conducted in triplicate.

### Animal vaccination

Animal experiments were conducted in accordance with the National Institutes of Health guidelines and the approval of Stanford Administrative Panel on Laboratory Animal Care (Protocol APLAC-32109). Six-to-seven week old female C57BL/6 (B6) mice were purchased from Charles River and housed in the animal facility at Stanford University. Mice were shaved on their right flank and received a subcutaneous injection of 100 µL of soluble or hydrogel vaccines under brief isoflurane anesthesia. Tail vein blood collection was performed according to the vaccination schedule.

A typical vaccine formulation dose consisted of RBD SARS-CoV-2 WT, BA.4/BA.5, SARS-CoV-1, MERS antigens (1:1:1:1 weight ratio, 10 μg total) and 3M-052/AF (1 μg), AddaVax (50 μL) or 3M-052/Alum (1 μg) as an adjuvant. Co-NTA liposomes were gently stirred with antigens for 30 mins at room temperature before being mixed with the polymer solution in PBS 1X. All soluble vaccines consisted of soluble antigens and adjuvants in PBS 1X. Formulations were loaded into a 26-gauge needle for injection.

#### Monovalent prime – bivalent boost SARS-CoV-2 RBD-LNH

Mice were first vaccinated with a soluble or hydrogel vaccine comprising SARS-CoV-2 WT RBD and 3M-052/AF, AddaVax or 3M-052/Alum as adjuvants (noted as Week 0). Mice were boosted on week 8 with a bivalent vaccine containing a 1:1 mass ratio of SARS-CoV-2 WT and BA.4/BA.5 RBDs adjuvanted as stated above (noted as Week 8).

#### Prime-boost tetravalent betacoronavirus RBD-LNH

Mice were primed and boosted with a soluble or hydrogel vaccine comprising a 1:1:1:1 mass ratio of SARS-CoV-2 WT, BA.4/BA.5, SARS-CoV-1 and MERS RBDs adjuvanted with 3M-052/AF, AddaVax or 3M-052/Alum (noted as Weeks 0 and 8).

#### Tetravalent betacoronavirus RBD-LNH in pre-exposed mice

Mice were primed and boosted with a soluble vaccine dose consisting of 1μg of Hexapro as the antigen and CpG/Alum as adjuvant (20 μg and 100 μg respectively, noted as Weeks 0 and 3). The same mice were vaccinated a third time on week 27 with a tetravalent betacoronavirus vaccine containing a 1:1:1:1 mass ratio of SARS-CoV-2 WT, BA.4/BA.5, SARS-CoV-1 and MERS RBDs adjuvanted with 3M-052/AF, 3M-052/Alum or AddaVax (noted as M6).

### Enzyme-linked immunosorbent assay (ELISA)

Serum anti-RBD IgG antibody endpoint titers were measured using an ELISA. Plates were washed 4 times between each step with a solution of PBS 1X containing 0.05% of Tween-20. Maxisorp plates were first coated with either SARS-CoV-2 Spike RBD, SARS-CoV-2 (Omicron BA.4/BA.5/BA.5.2) Spike RBD, SARS-CoV Spike/RBD or MERS-CoV Spike/RBD proteins at 2 µg/mL in PBS 1X overnight at 4 °C and subsequently blocked with PBS 1X containing 1 wt% BSA for 1 h at 25 °C. The sera were serially diluted starting at 1:100 or 1:200 followed by a 4-fold dilution and incubated in the coated plates for 2 h at 25 °C. Goat-anti-mouse IgG Fc-HRP (1:10,000) was added for 1 h at 25 °C and plates were developed with TMB substrate for 6 mins.

The reaction was stopped with 1 M aqueous HCl solution and a colorimetric readout was performed at 450 nm in absorbance using a Synergy H1 microplate reader (BioTek Instruments). Absorbance versus dilution curves of each dataset were generated and fitted with a five-parameter non-linear logistic regression using GraphPad 10. The dilution titer value at which the endpoint threshold of 0.1 was crossed for each curve was reported as IgG endpoint titer. Samples failing to meet endpoint threshold at a 1:100 dilution were set to below the limit quantitation (1:25 dilution). Breadth experiments included SARS-CoV-2 Spike, SARS-CoV-2 (Delta B.1.617.2) Spike S1, SARS-CoV-2 (Omicron B.1.1.529) Spike S1, XBB.1.5 Omicron Spike RBD, SARS-CoV Spike RBD, and SARS-CoV Spike RBD.

IgG isotypes were measured according to the same protocol as previously stated. Goat anti-mouse IgG1 and IgG2c Fc-HRP secondary antibodies were added at a dilution of 1:20,000 and an endpoint threshold of 0.2 was used to report titer values.

The power law decay model was used to estimate the decay half-life of binding antibody endpoint titers. The equation dAb/dt = −k/t × Ab and Ab = C × t^(−k) was fitted using Matlab’s fitnlm function (R2024a, Mathworks) to the longitudinal data starting from D14 after the initial immunization. Ab is the RBD-specific antibody-binding endpoint titers and k is the power law decay rate. Longitudinal data from three time points were used to estimate the decay rate and the corresponding half-lives were calculated as t1/2 = 0.5^(1/k).

### Enzyme-linked immunosorbent spot (ELISpot)

Single color mouse IgG ELISPOT kit was purchased from Cellular Technology Limited. Bone marrow cells were harvested 2 weeks post-boost in the respective vaccination regime following immunization schedules outline in Figure 2 and Figure 5. Wells were first coated with 80 µL of either SARS-CoV-2 Spike RBD, SARS-CoV-2 (Omicron BA.4/BA.5/BA.5.2) Spike RBD, SARS-CoV Spike/RBD or MERS-CoV Spike/RBD proteins at 25 µg/mL in PBS 1X or Capture Solution prepared according to the manufacturer’s specifications overnight at 4 °C and subsequently blocked with assay medium (RPMI 1640 with 10% FBS and 2mM L-glutamine). 300,000 bone marrow cells per sample were pipetted into the wells and spots were developed following manufacturer’s instruction (Immunospot S6 Ultra M2).

### Neutralization assays using pseudotyped lentivirus

Spike-pseudotyped lentiviruses encoding a luciferase-ZsGreen reporter were produced using a five-plasmid system as previously described.^74^ Briefly, the plasmid system uses the packaging vector pHAGE-Luc2-IRES-ZsGreen(NR-52516), three helper plasmids HDM-Hgpm2 (NR-52517), HDM-tat1b (NR-52518), pRC-CMV-Rev1b (NR-52519), and a plasmid encoding the viral spike protein. The wild-type SARS-CoV-2 spike plasmid (HDM-SARS2-spike-delta21, Addgene #155130) was cloned with the additional D614G substitution. The Omicron BA.4/5 spike plasmid contained the additional substitutions T19I, Δ24-26, A27S, G142D, V213G, G339D, S371F, S373P, S375F, T376A, D405N, R408S, K417N, N440K, L452R, S477N, T478K, E484A, F486V, Q493R, Q498R, N501Y, Y505H, H655Y, N679K, P681H, N764K, D796Y, Q954H, N969K (sequence ID: UFO69279.1). One day prior to transfection, Expi-293F cells were diluted to 3×10^6^ cells/mL in 200 ml. Transfection mixture was prepared by adding 200 µg of packaging vector, 68 µg of spike plasmid, and 44 µg of helper plasmids to 20 mL of FreeStyle293/Expi-293 media followed by the dropwise addition and mixing of 600 µL BioT. After a 10 min incubation at room temperature, the transfection mixture was added to the Expi-293F cells after which cells were boosted with D-glucose (4 g/L) and valproic acid (3 mM). After 72 h, the cells were spun down, the culture supernatant was collected and filtered through a 0.45-µm filter, and the virus was aliquoted, flash frozen in liquid nitrogen, and stored at −80 °C. All pseudotyped lentiviruses were titrated in HEK-293T cells prior to usage.

One day prior to infection, HeLa cells stably overexpressing human angiotensin-converting enzyme 2 (ACE2) and transmembrane serine protease 2 (TMPRSS2) were seeded at a density of 1×10^4^ cells/well in white-walled 96-well plates (ThermoFisher Scientific). The following day, heat-inactivated mouse antisera (56 °C, 30 mins) was serially diluted in D10 media and mixed with pseudotyped lentivirus diluted in D10 media supplemented with polybrene (Sigma-Aldrich) at a final concentration of 5 µg/mL. After 1 h incubation at 37 °C, 100 µL of the antisera/viral dilutions were transferred to wells containing the seeded HeLa/ACE2/TMPRSS2 cells. After 48 h, the medium was removed from the wells, and cells were lysed by the addition of 100 µL of luciferase substrates (BriteLite Plus, Perkin Elmer). Luminescent signals were recorded on a microplate reader (BioTek SynergyTM HT or Tecan M200) after shaking for 30 sec. Infection percent was normalized to the signal in cells only wells (0 % infection) and virus only wells (100 % infection) on each plate. The neutralization titer (NT_50_) was defined as the serum dilution concentration at which 50 % virus infection was measured. Values were plotted and fitted in GraphPad Prism.

### Flow Cytometry of draining lymph nodes

Mice were euthanized using carbon dioxide 14 days post immunization to collect inguinal lymph nodes. Following their dissociation into single cell suspensions, staining for viability was performed using Ghost Dye Violet 510 (Tonbo Biosciences, cat. Number 13-0870-T100) for 5 mins on ice and washed with FACS buffer (PBS 1X with 2% FBS, 1mM EDTA). Then, Fc receptors were blocked using anti-CD16/CD38 antibody (clone: 2.4G2, BD Biosciences, cat. Number 553142) for 5 mins on ice and stained with fluorochrome conjugated antibodies: CD19 (PerCP-Cy5.5, clone: 1D3, BioLegend, cat. Number 152406), CD95 (PE-Cy7, clone: Jo2, BD Biosciences, cat. Number 557653), GL7 (AF488, clone: GL7, BioLegend, cat. Number 144613), CD3 (AF700, clone: 17A2, BioLegend, cat. Number 100216), CD4 (BV650, clone: GK1.5, BioLegend, cat. Number 100469), CXCR5 (BV711, clone: L138D7, BioLegend, cat. Number 145529), PD1 (PE-Dazzle^TM^594, clone: 29F.1A12, BioLegend, cat. Number 135228), anti-spike tetramer (AF647) and anti-spike tetramer (Cy3) for 30 mins on ice. Tetramers were prepared on ice by adding AF647-or Cy3-Streptavidin (Thermo Scientific, cat. Number S21374; ThermoFisher Scientific, cat. Number 434315) to 7.1 μM biotinylated WT SARS-CoV-2 spike trimer (Sino Biological, cat. Number 40589-V27B-B) in five steps, once every 20 mins, for a final molar ratio of 5:1 spike protein to dye and concentration of 0.8 μM. Cells were washed and fixed with 4% PFA on ice. After a washing step, cells were analyzed on an LSRII flow cytometer (BD Biosciences). Data were analyzed with FlowJ 10 (FlowJo, LLC).

### Flow cytometry of LNH immune infiltrate

Mice were shaved and injected subcutaneously on the right flank with 100 μL of empty or vaccine LNHs loaded with WT SARS-CoV-2 RBD protein (10 μg per dose) and 3M-052/AF adjuvant (1 μg per dose). Three days after injection, mice were euthanized using carbon dioxide. Hydrogel depots were extracted and placed in microcentrifuge tubes with 750 μL of FACS buffer (PBS 1X, 3% heat inactivated FBS, 1 mM EDTA). Hydrogels were mechanically disrupted to single cell suspensions using Kimble BioMasherIIs (DWK Life Sciences). Suspensions were passed through 70 μm cell filters (Celltreat, 229484) into 15 mL Falcon tubes, spun at 400 rcf for 4 mins, resuspended in 300 μL of PBS 1X, and counted using acridine orange/propidium iodide cell viability stain (Vitascientific, LGBD10012) with a Luna-FL dual Fluorescence cell counter (Logos biosystems). One million live cells per sample were transferred to a 96-well conical bottom plate (Thermo Scientific, 249570) and stained using the following protocol. Hydrogel samples were first stained with 100 μL of Live/Dead Fixable Near-IR (Thermo Scientific, L34975) for 30 mins at room temperature, quenched with 100 μL of FACS buffer, and spun at 935 rcf for 2 mins. Samples were then incubated with 50 μL of anti-mouse CD16/CD32 (1:50 dilution; BD, 553142) for 5 mins on ice before incubating with 50 μL of full antibody stain for 30 mins on ice. Samples were quenched with 100 μL of FACS buffer, spun at 935 rcf for 2 mins, and resuspended in 60 μL of FACS buffer. Samples were run on the BD FACSymphony A5 SORP in the Stanford Shared FACS Facility and data were analyzed in FlowJo.

Hydrogel full antibody stain included anti-I-A/I-E (1:800 dilution; FITC; BioLegend, 107605), anti-CD45 (1:800; AF700; BioLegend, 103127), anti-Ly6C (1:400; BV570; BioLegend, 128029), anti-XCR1 (1:200; AF647; BioLegend, 148213), anti-F4/80 (1:200; BV421; BioLegend, 123137), anti-Ly6G (1:200; BV711; BioLegend, 127643), anti-CD3e (1:200; PerCP-eFluor710; Thermo Scientific, 46-0033-82), anti-CD19 (1:200; PE-Cy7; BioLegend, 115519), anti-CD11c (1:200; PE; BioLegend, 117307), anti-CD11b (1:100; BV510; Fisher Scientific, 50-112-9846), and anti-NK1.1 (1:100; BV605; BioLegend, 108753).

### Statistical analyses

Animals were caged blocked for all in vivo experiments except for the cellular infiltration study. A sample size of n=4-5 was used and data are presented as mean +/− standard error of mean (SEM) or median +/− interquartile range as specified in the corresponding figure captions. Comparisons between the treatment of interest (vaccine-loaded hydrogel) and each control (empty hydrogel or soluble vaccines) were assessed using the general linear model followed by Student’s t-test in JMP and accounted for cage blocking. Comparison between multiple groups were conducted similarly, using the general linear model (GLM) followed by Tukey HSD procedure test in JMP. Data were used raw or transformed (log10) when necessary to result in the highest coefficient of determination and best fit the model. To meet statistical requirements for normalized variance, cell infiltration data that is presented as percentages was transformed using the following equation: y = ln(x/(100-x)), x being the original data and y the transformed value used in statistical analysis. Statistical significance was considered as the following: * for *p* < 0.05, ** for *p* < 0.01 and *** for *p* < 0.001. Selected *p* values are shown in the figures and all *p* values are in the supporting information.

## Supplemental information description

Document S1. Figures S1-S10, Tables S1-S18.

## Acknowledgements

This work was financially supported by the Bill & Melinda Gates Foundation (INV-027411). J.B. is thankful for a Marie-Curie fellowship from the European Union (H2020; No. 101030481). B.S.O. is grateful for an Eastman Kodak Fellowship. E.L.M. was supported by the NIH Biotechnology Training Program (T32 GM008412). J.Y. and C.K.J. are thankful for a National Science Foundation Graduate Research Fellowship Program. The authors extend their gratitude to the members of the Appel lab for their scientific expertise and help, specifically Dr. Olivia M. Saouaf and Ye E. Song. The authors also thank Jodi Hanson from CTL for her consultation on ELISpot. Flow cytometry was performed on an instrument in the Stanford Shared FACS Facility obtained using NIH S10 Shared Instrument Grant (1S10OD026831-01).

## Authors contribution

Conceptualization, J.B., J.H.K., and E.A.A.; methodology, J.B., J.H.K., E.L.M., B.S.O., T.U.J.B, and A.U.; formal analysis, J.B., J.H.K., E.L.M., and B.S.O.; investigation, J.B., J.H.K., E.L.M., B.S.O., J.Y., T.U.J.B, A.U., and C.K.J.; visualization, J.B., and J.H.K.; writing – original draft, J.B., and J.H.K.; writing – review & editing, J.B., J.H.K., B.S.O., E.L.M., S.L. and E.A.A.; funding acquisition, J.B., S.L., and E.A.A.; supervision, E.A.A.

