## Supporting Information for "Sustained exposure to multivalent antigen-decorated nanoparticles generates broad anti-coronavirus responses"

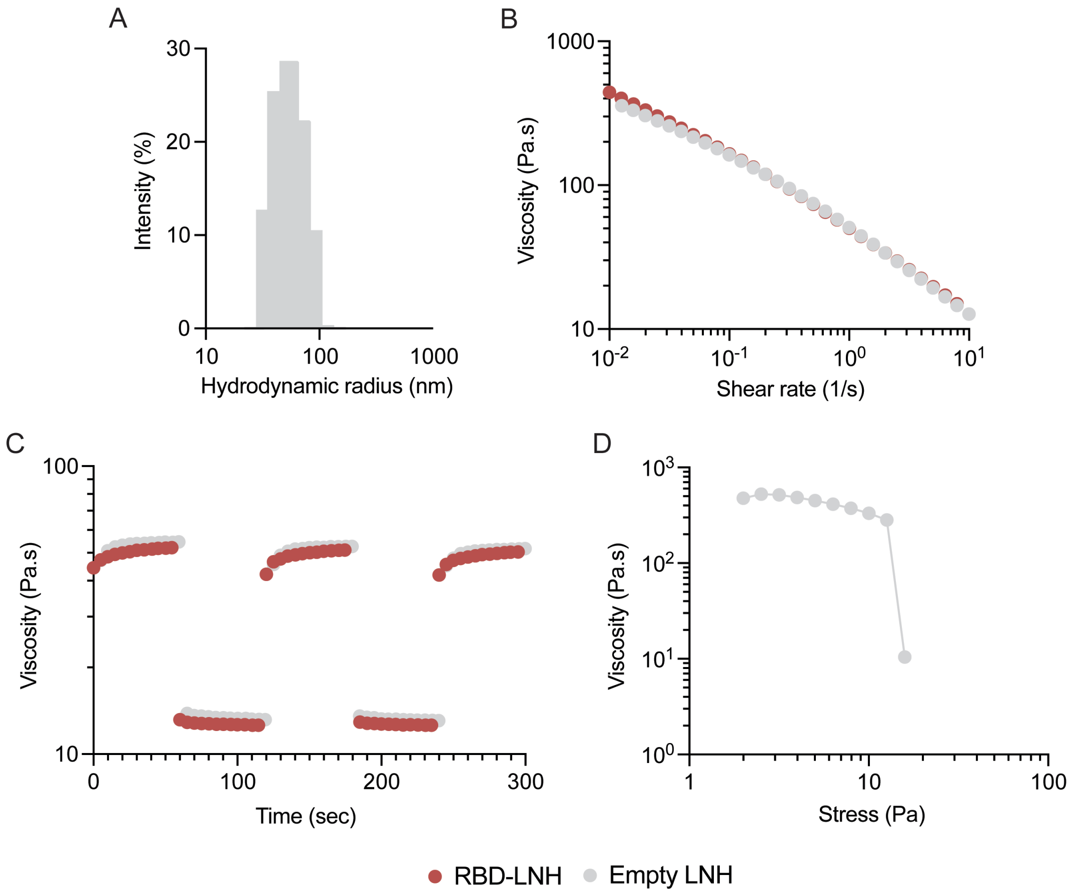

**Figure S1. Characterization of liposomal hydrogels. a.** Distribution of the hydrodynamic radii of plain liposomes measured by Dynamic Light Scattering. **b.** Shear-dependent flow rheology of empty and vaccine-loaded LNHs. c. Step-shear measurements of empty and vaccine-loaded LNHs over cycles of alternating high shear (10 $s^{-1}$) and low shear (0.1 $s^{-1}$) rates. d. Stress-controlled yield stress measurement of empty LNH.

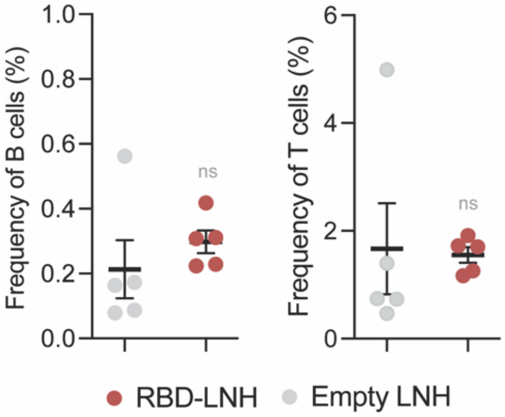

**Figure S2. Cellular infiltration in liposomal hydrogels.** Frequency of B cells and T cells infiltrating empty (grey) and vaccine-loaded (ref) LNHs (WT SARS-CoV-2 RBD, 3M-052/AF) 3 days after subcutaneous injection. Data are shown as mean +/- SEM (n=5). *p* values were calculated from the general linear model followed by Student's t-test and reported in Table S1.

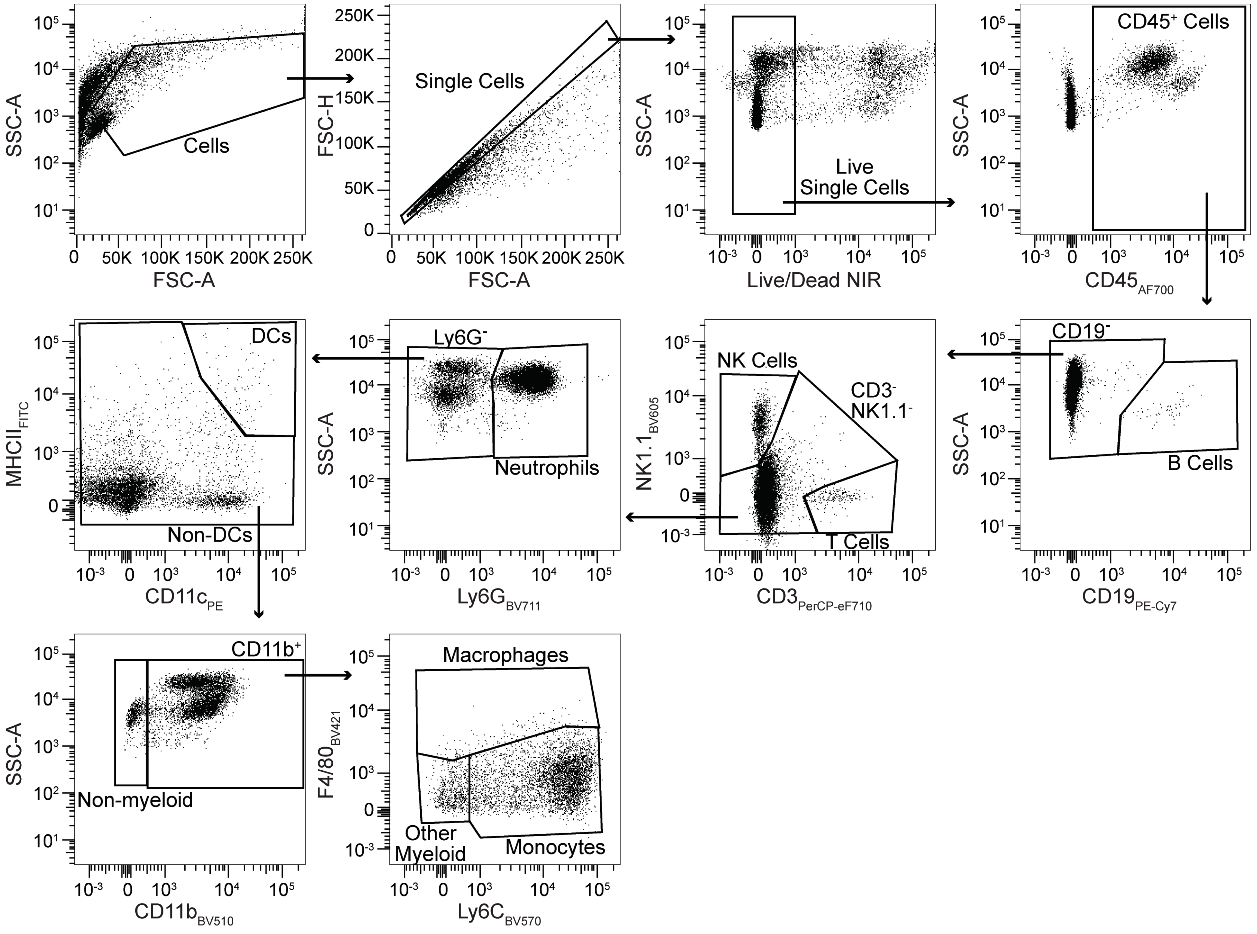

**Figure S3. Representative gating strategy for cellular infiltration.**

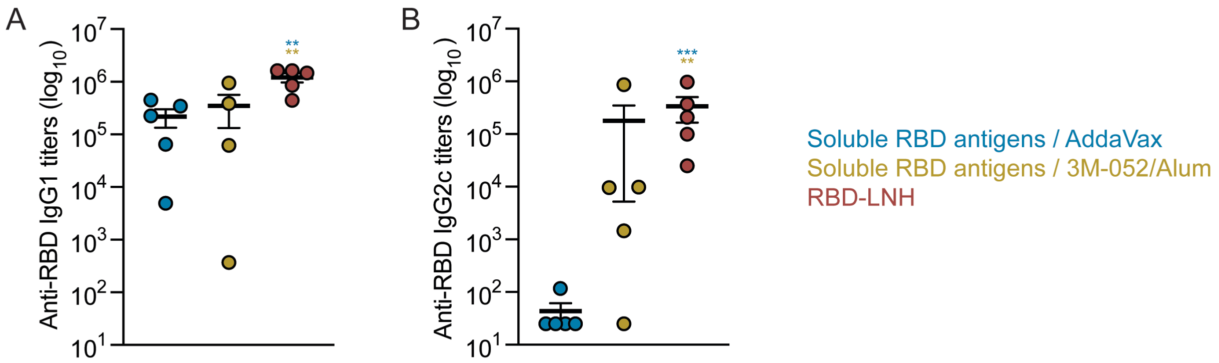

**Figure S4. Antibody subtype response to a monovalent prime + bivalent boost SARS-CoV-2 vaccination.** Anti-RBD IgG1 and IgG2c titers from mice sera collected on Week 5, 2 weeks after boost. Data are shown as mean +/- SEM (n=5). *p* values were calculated from the general linear model followed by Student's t-test and reported in Table S4.

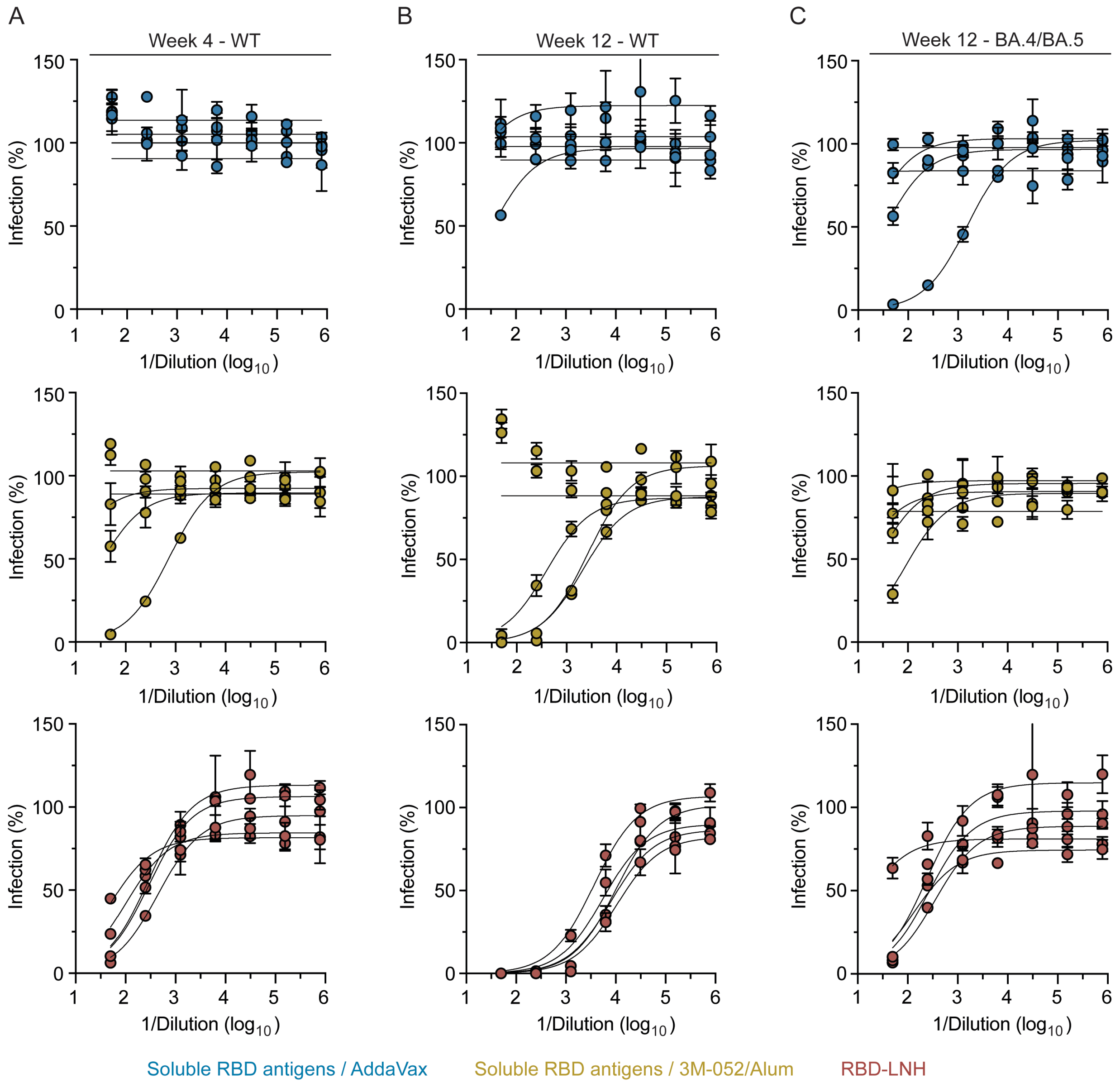

**Figure S5. Neutralizing antibodies activity in mice vaccinated with a monovalent prime + bivalent boost SARS-CoV-2 vaccine.** Percent infectivity of the neutralizing antibodies from a range of dilution of mice sera vaccinated with RBD-LNH adjuvanted with 3M-052/AF or the two soluble controls adjuvanted with AddaVax or 3M-052/Alum against SARS-CoV-2 WT and BA.4/BA.5 spike-pseudotyped lentiviruses a. 4 weeks post-prime (Week 4, WT) and b.c. 4 weeks post boost (Week 12, WT/BA.4/BA.5).

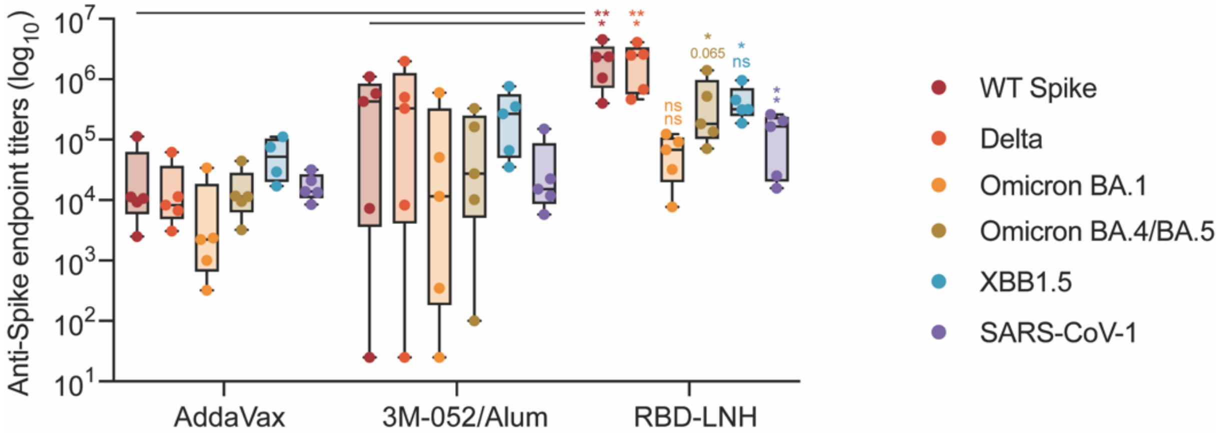

**Figure S6. Breadth of antibodies induced by monovalent prime + bivalent boost SARS-CoV-2 vaccine.** Anti-Spike IgG endpoint titers measured for SARS-CoV-2 Omicron BA.1, XBB1.5, BA.4/BA.5, Delta, WT and SARS-CoV-1 4 weeks post boost (W12). Data are shown as mean +/- SEM (n = 5). *p* values were calculated from the general linear model followed by Student's t-test and reported in Table S6.

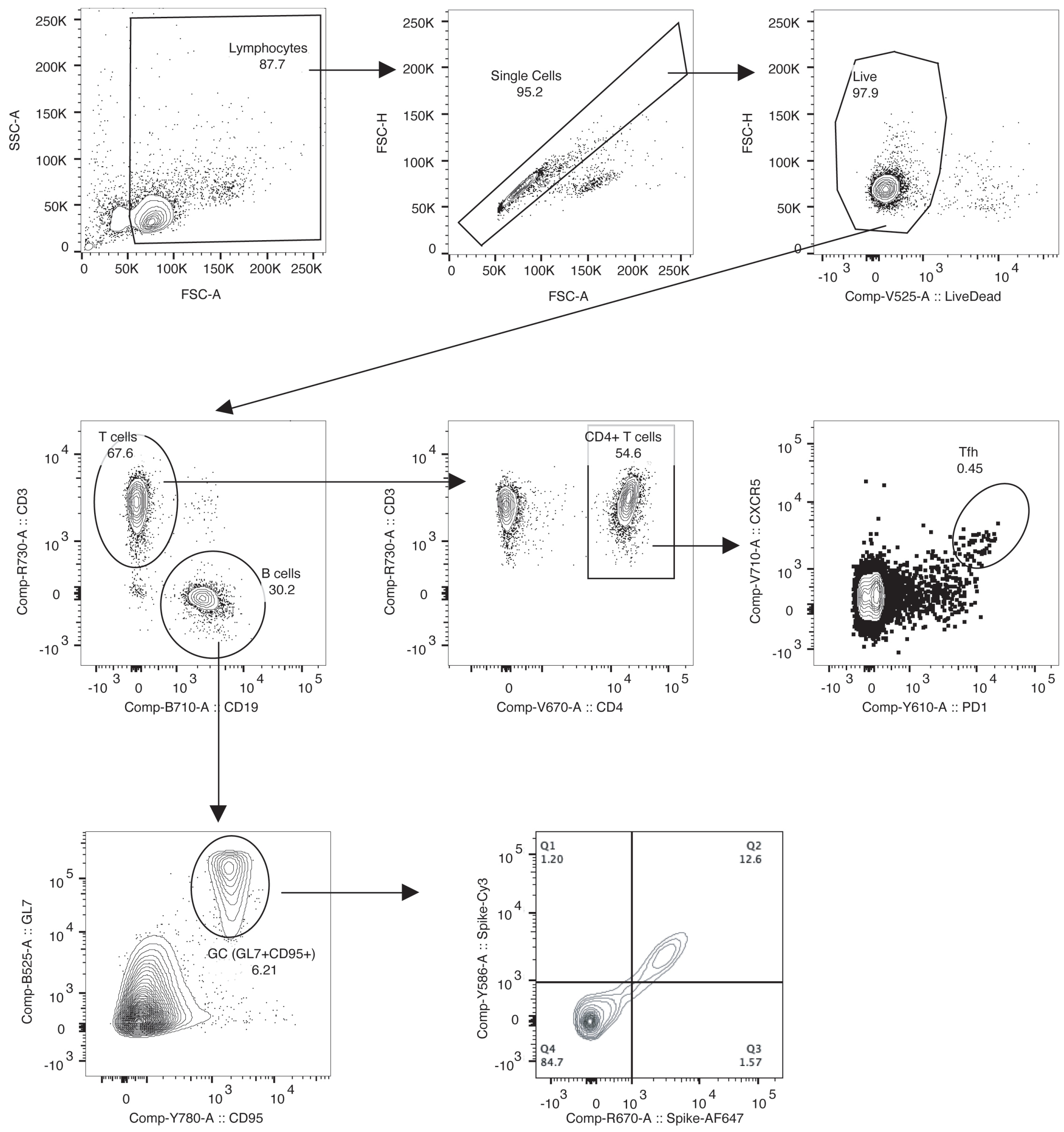

**Figure S7. Representative gating strategy for germinal center (GC) analysis.**

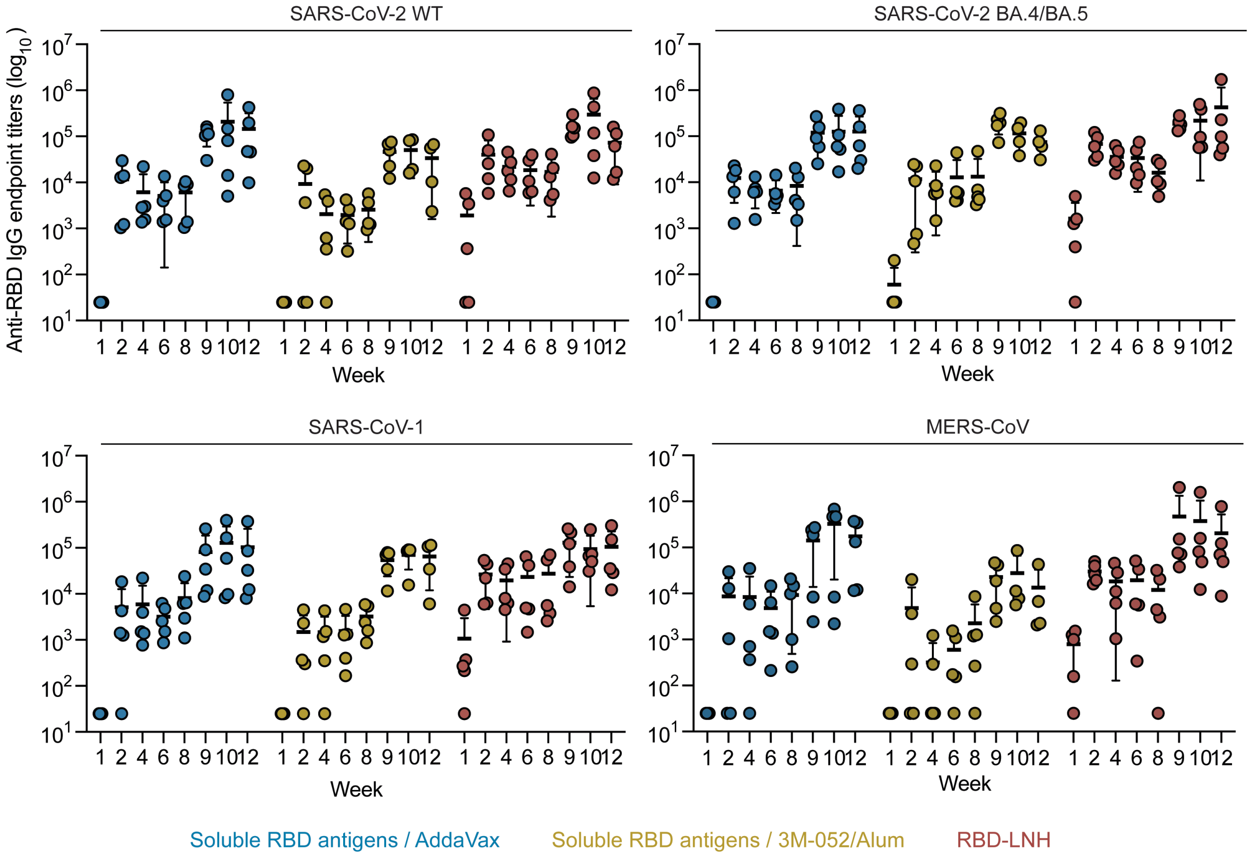

**Figure S8. Humoral antibody response following a prime-boost tetravalent betacoronavirus vaccination.** Anti-RBD IgG antibody endpoint titers measured over 12 weeks against SARS-CoV-2 WT, BA.4/BA.5, SARS-CoV-1 and MERS-CoV strains from mice immunized with tetravalent RBD-LNH adjuvanted with 3M-052/AF or the two soluble controls adjuvanted with AddaVax or 3M-052/Alum. Data are shown as mean +/- SEM (n = 4-5).

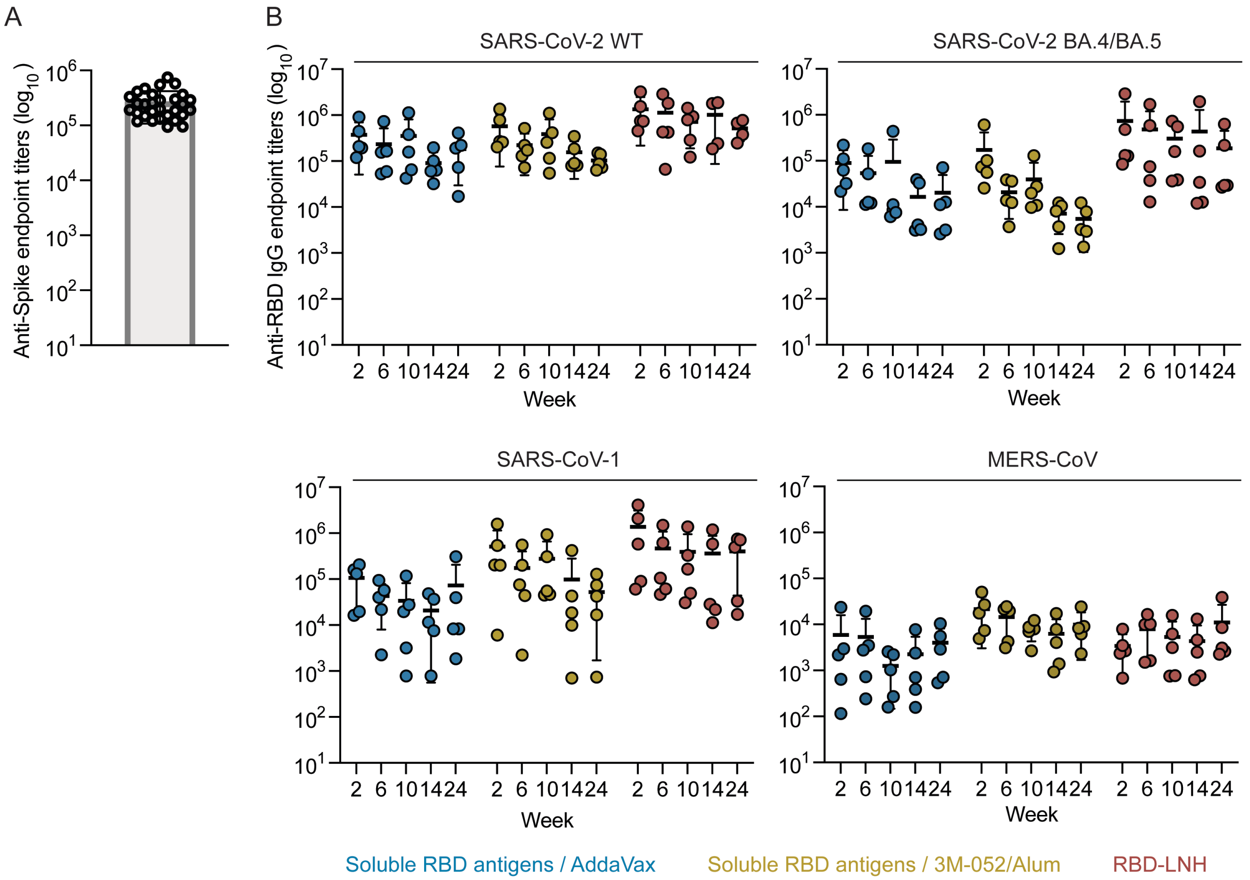

**Figure S9. Humoral antibody response of pre-exposed mice boosted with tetravalent coronavirus vaccine. a.** Anti-SARS-CoV-2 Spike IgG antibody endpoint titers measured six months following a prime-boost immunization of soluble Hexapro adjuvanted with CpG/Alum. Data are shown as mean +/- SEM (n = 30). **b.** Anti-RBD IgG antibody endpoint titers measured over 24 weeks against SARS-CoV-2 WT, BA.4/BA.5, SARS-CoV-1 and MERS-CoV strains from pre-exposed mice boosted with tetravalent RBD-LNH adjuvanted with 3M-052/AF or the two soluble controls adjuvanted with AddaVax or 3M-052/Alum. Weeks 1 and 24 refer to W29 and M12 in Figure 7. Data are shown as mean +/- SEM (n = 4-5).

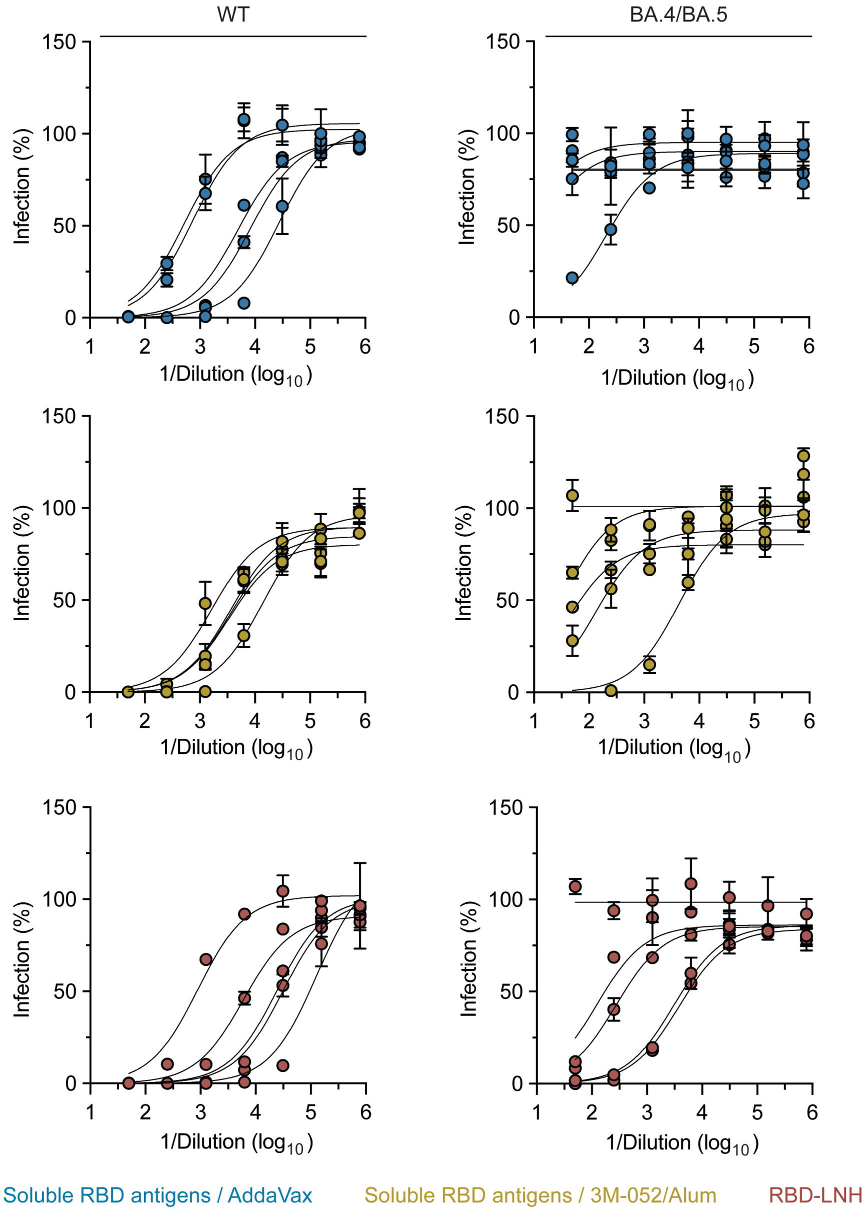

**Figure S10. Neutralizing antibodies activity in pre-exposed mice boosted with tetravalent coronavirus vaccine.** Percent infectivity of the neutralizing antibodies from a range of dilution of mice sera vaccinated with RBD-LNH adjuvanted with 3M-052/AF or the two soluble controls adjuvanted with AddaVax or 3M-052/Alum against SARS-CoV-2 WT and BA.4/BA.5 spike-pseudotyped lentiviruses 14 weeks post-boost.

**Table S1.** *p* values for the frequency of cells infiltrating empty and vaccine loaded LNH (general linear model followed by Student's t-test).

| | | Adjusted *p* values | |
|---|---|---|---|
| Frequency of macrophages | Empty vs. Vaccine-loaded | 0.2673 | ns |
| Frequency of monocytes | Empty vs. Vaccine-loaded | 0.1100 | ns |
| Frequency of B cells | Empty vs. Vaccine-loaded | 0.4032 | ns |
| Frequency of T cells | Empty vs. Vaccine-loaded | 0.8981 | ns |
| Frequency of NK cells | Empty vs. Vaccine-loaded | 0.0037 | ** |

**Table S2.** *p* values for anti-RBD endpoint IgG titers elicited by a SARS-CoV-2 monovalent prime + bivalent boost of different vaccine formulations (general linear model followed by Student's t-test).

| | | Adjusted *p* values | |
|---|---|---|---|
| Anti-WT RBD IgG titers ($log_{10}$) Week 2 | AddaVax vs. RBD-LNH | 0.0085 | ** |
| | 3M-052/Alum vs. RBD-LNH | 0.0202 | * |
| Anti-WT RBD IgG titers ($log_{10}$) Week 10 | AddaVax vs. RBD-LNH | 0.0003 | *** |
| | 3M-052/Alum vs. RBD-LNH | 0.0031 | ** |
| Anti-WT RBD IgG titers ($log_{10}$) Week 12 | AddaVax vs. RBD-LNH | 0.0703 | ns |
| | 3M-052/Alum vs. RBD-LNH | 0.0863 | ns |

| | | Adjusted *p* values | |
|---|---|---|---|
| Anti-BA.4/BA.5 RBD IgG titers ($log_{10}$) Week 2 | AddaVax vs. RBD-LNH | 0.9948 | ns |
| | 3M-052/Alum vs. RBD-LNH | 0.6733 | ns |
| Anti-BA.4/BA.5 RBD IgG titers ($log_{10}$) Week 11 | AddaVax vs. RBD-LNH | 0.0526 | ns |
| | 3M-052/Alum vs. RBD-LNH | 0.0531 | ns |
| Anti-BA.4/BA.5 RBD IgG titers ($log_{10}$) Week 12 | AddaVax vs. RBD-LNH | 0.0559 | ns |
| | 3M-052/Alum vs. RBD-LNH | 0.0841 | ns |

**Table S3.** *p* values for the area under the curve (AUC) over 8 (post-prime) and 12 (post-boost) weeks of anti-RBD endpoint IgG titers elicited by a SARS-CoV-2 monovalent prime + bivalent boost of different vaccine formulations (general linear model followed by Student's t-test).

| | | Adjusted *p* values | |
|---|---|---|---|
| Anti-WT RBD IgG titers ($log_{10}$) AUC (post-prime) | AddaVax vs. RBD-LNH | 0.0036 | ** |
| | 3M-052/Alum vs. RBD-LNH | 0.0959 | ns |
| Anti-WT RBD IgG titers ($log_{10}$) AUC (post-boost) | AddaVax vs. RBD-LNH | 0.0005 | *** |
| | 3M-052/Alum vs. RBD-LNH | 0.0026 | ** |

| | | Adjusted *p* values | |
|---|---|---|---|
| Anti-BA.4/BA.5 RBD IgG titers ($log_{10}$) AUC (post-prime) | AddaVax vs. RBD-LNH | 0.1211 | ns |
| | 3M-052/Alum vs. RBD-LNH | 0.4641 | ns |
| Anti-BA.4/BA.5 RBD IgG titers ($log_{10}$) AUC (post-boost) | AddaVax vs. RBD-LNH | 0.0088 | ** |
| | 3M-052/Alum vs. RBD-LNH | 0.0239 | * |

**Table S4.** *p* values for anti-RBD endpoint IgG1 and IgG2c titers elicited by a SARS-CoV-2 monovalent prime - bivalent boost of different vaccine formulations (general linear model followed by Student's t-test).

| | | Adjusted *p* values | |
|---|---|---|---|
| Anti-WT RBD IgG1 titers ($log_{10}$) Week 10 | AddaVax vs. RBD-LNH | 0.0020 | ** |
| | 3M-052/Alum vs. RBD-LNH | 0.0053 | ** |
| Anti-WT RBD IgG2c titers ($log_{10}$) Week 10 | AddaVax vs. RBD-LNH | 0.0002 | *** |
| | 3M-052/Alum vs. RBD-LNH | 0.0058 | ** |

**Table S5.** *p* values for $NT_{50}$ neutralization titers elicited by a SARS-CoV-2 monovalent prime + bivalent boost of different vaccine formulations (general linear model followed by Student's t-test).

| | | Adjusted *p* values | |
|---|---|---|---|
| Neutralization $NT_{50}$ titers ($log_{10}$) WT - Week 4 | AddaVax vs. RBD-LNH | 0.0262 | * |
| | 3M-052/Alum vs. RBD-LNH | 0.1540 | ns |
| Neutralization $NT_{50}$ titers ($log_{10}$) WT - Week 12 | AddaVax vs. RBD-LNH | <0.0001 | *** |
| | 3M-052/Alum vs. RBD-LNH | 0.0025 | ** |
| Neutralization $NT_{50}$ titers ($log_{10}$) Omicron BA.4/BA.5/BA.5.2 - Week 12 | AddaVax vs. RBD-LNH | 0.3581 | ns |
| | 3M-052/Alum vs. RBD-LNH | 0.5478 | ns |

**Table S6.** *p* values for anti-spike endpoint titers of SARS-related strains elicited by a SARS-CoV-2 monovalent prime + bivalent boost of different vaccine formulations (general linear model followed by Student's t-test).

| | | Adjusted *p* values | |
|---|---|---|---|
| IgG titers ($log_{10}$) - WT spike Week 12 | AddaVax vs. RBD-LNH | 0.0053 | ** |
| | 3M-052/Alum vs. RBD-LNH | 0.0149 | * |
| IgG titers ($log_{10}$) - Delta B.1.617.2 Week 12 | AddaVax vs. RBD-LNH | 0.0042 | ** |
| | 3M-052/Alum vs. RBD-LNH | 0.0202 | * |
| IgG titers ($log_{10}$) - Omicron B.1.1.529 Week 12 | AddaVax vs. RBD-LNH | 0.5675 | ns |
| | 3M-052/Alum vs. RBD-LNH | 0.4931 | ns |
| IgG titers ($log_{10}$) - Omicron BA.4/BA.5/BA.5.2 Week 12 | AddaVax vs. RBD-LNH | 0.0389 | * |
| | 3M-052/Alum vs. RBD-LNH | 0.0650 | ns |
| IgG titers ($log_{10}$) - Omicron XBB1.5 Week 12 | AddaVax vs. RBD-LNH | 0.0202 | * |
| | 3M-052/Alum vs. RBD-LNH | 0.2938 | ns |
| IgG titers ($log_{10}$) - SARS-CoV-1 Week 12 | AddaVax vs. RBD-LNH | 0.0111 | * |
| | 3M-052/Alum vs. RBD-LNH | 0.0234 | * |

**Table S7.** *p* values for total GCBC, antigen-specific GCBC and Tfh counts elicited by a SARS-CoV-2 monovalent prime + bivalent boost of different vaccine formulations (general linear model followed by Student's t-test).

| | | Adjusted *p* values | |
|---|---|---|---|
| Total GCBCs ($log_{10}$) Week 2 | AddaVax vs. RBD-LNH | 0.1755 | ns |
| | 3M-052/Alum vs. RBD-LNH | 0.5499 | ns |
| WT-specific GCBCs ($log_{10}$) Week 2 | AddaVax vs. RBD-LNH | 0.6938 | ns |
| | 3M-052/Alum vs. RBD-LNH | 0.3098 | ns |
| Total Tfh cells ($log_{10}$) Week 2 | AddaVax vs. RBD-LNH | 0.0263 | * |
| | 3M-052/Alum vs. RBD-LNH | 0.0500 | ns |

**Table S8.** *p* values for percent of WT and BA.4/BA.5 antigen secreting cells (ASCs) isolated from bone marrow, elicited by a SARS-CoV-2 monovalent prime + bivalent boost of different vaccine formulations (general linear model followed by Student's t-test).

| | | Adjusted *p* values | |
|---|---|---|---|
| $WT^{+}$ ASCs Week 2 | AddaVax vs. RBD-LNH | 0.8146 | ns |
| | 3M-052/Alum vs. RBD-LNH | 0.9604 | ns |
| $BA.4/BA.5^{+}$ ASCs Week 2 | AddaVax vs. RBD-LNH | 0.8618 | ns |
| | 3M-052/Alum vs. RBD-LNH | 0.5720 | ns |

**Table S9.** *p* values for anti-RBD endpoint IgG titers elicited by a tetravalent betacoronavirus prime-boost of different vaccine formulations (general linear model followed by Student's t-test).

| | | Adjusted *p* values | |
|---|---|---|---|
| Anti-WT RBD IgG titers ($log_{10}$) Week 2 | AddaVax vs. RBD-LNH | 0.3772 | ns |
| | 3M-052/Alum vs. RBD-LNH | 0.0726 | ns |
| Anti-WT RBD IgG titers ($log_{10}$) Week 10 | AddaVax vs. RBD-LNH | 0.2134 | ns |
| | 3M-052/Alum vs. RBD-LNH | 0.0349 | * |

| | | Adjusted *p* values | |
|---|---|---|---|
| Anti-BA.4/BA.5 RBD IgG titers ($log_{10}$) Week 2 | AddaVax vs. RBD-LNH | 0.0049 | ** |
| | 3M-052/Alum vs. RBD-LNH | 0.0046 | ** |
| Anti-BA.4/BA.5 RBD IgG titers ($log_{10}$) Week 10 | AddaVax vs. RBD-LNH | 0.0475 | * |
| | 3M-052/Alum vs. RBD-LNH | 0.1159 | ns |

| | | Adjusted *p* values | |
|---|---|---|---|
| Anti-SARS-CoV-1 RBD IgG titers ($log_{10}$) Week 2 | AddaVax vs. RBD-LNH | 0.0644 | ns |
| | 3M-052/Alum vs. RBD-LNH | 0.0192 | * |
| Anti-SARS-CoV-1 RBD IgG titers ($log_{10}$) Week 10 | AddaVax vs. RBD-LNH | 0.6421 | ns |
| | 3M-052/Alum vs. RBD-LNH | 0.6026 | ns |

| | | Adjusted *p* values | |
|---|---|---|---|
| Anti-MERS-CoV RBD IgG titers ($log_{10}$) Week 2 | AddaVax vs. RBD-LNH | 0.0365 | * |
| | 3M-052/Alum vs. RBD-LNH | 0.0180 | * |
| Anti-MERS-CoV RBD IgG titers ($log_{10}$) Week 10 | AddaVax vs. RBD-LNH | 0.8056 | ns |
| | 3M-052/Alum vs. RBD-LNH | 0.0951 | ns |

**Table S10.** *p* values for the area under the curve (AUC) over 8 (post-prime) and 12 (post-boost) weeks of anti-RBD endpoint IgG titers elicited by a tetravalent betacoronavirus prime-boost of different vaccine formulations (general linear model followed by Student's t-test).

| | | Adjusted *p* values | |
|---|---|---|---|
| Anti-WT RBD IgG titers ($log_{10}$) AUC (post-prime) | AddaVax vs. RBD-LNH | 0.0518 | ns |
| | 3M-052/Alum vs. RBD-LNH | 0.0261 | * |
| Anti-WT RBD IgG titers ($log_{10}$) AUC (post-boost) | AddaVax vs. RBD-LNH | 0.3079 | ns |
| | 3M-052/Alum vs. RBD-LNH | 0.0315 | * |

| | | Adjusted *p* values | |
|---|---|---|---|
| Anti-BA.4/BA.5 RBD IgG titers ($log_{10}$) AUC (post-prime) | AddaVax vs. RBD-LNH | 0.0035 | ** |
| | 3M-052/Alum vs. RBD-LNH | 0.0039 | ** |
| Anti-BA.4/BA.5 RBD IgG titers ($log_{10}$) AUC (post-boost) | AddaVax vs. RBD-LNH | 0.0109 | * |
| | 3M-052/Alum vs. RBD-LNH | 0.0389 | * |

| | | Adjusted *p* values | |
|---|---|---|---|
| Anti-SARS-CoV-1 RBD IgG titers ($log_{10}$) AUC (post-prime) | AddaVax vs. RBD-LNH | 0.0881 | ns |
| | 3M-052/Alum vs. RBD-LNH | 0.0127 | * |
| Anti-SARS-CoV-1 RBD IgG titers ($log_{10}$) AUC (post-boost) | AddaVax vs. RBD-LNH | 0.6684 | ns |
| | 3M-052/Alum vs. RBD-LNH | 0.3644 | ns |

| | | Adjusted *p* values | |
|---|---|---|---|
| Anti-MERS-CoV RBD IgG titers ($log_{10}$) AUC (post-prime) | AddaVax vs. RBD-LNH | 0.0934 | ns |
| | 3M-052/Alum vs. RBD-LNH | 0.0250 | * |
| Anti-MERS-CoV RBD IgG titers ($log_{10}$) AUC (post-boost) | AddaVax vs. RBD-LNH | 0.5624 | ns |
| | 3M-052/Alum vs. RBD-LNH | 0.0542 | ns |

**Table S11.** *p* values for the area under the curve (AUC) over 12 weeks (post-boost) comparing anti-RBD endpoint IgG titers elicited by a tetravalent betacoronavirus prime-boost vaccine formulation across antigens (general linear model followed by Tukey's HSD multiple comparisons).

| | | Adjusted *p* values |
|---|---|---|
| AUC post-boost RBD IgG titers ($log_{10}$) Soluble AddaVax | WT vs. BA.4/BA.5 | 0.8925 |
| | WT vs. SARS-CoV-1 | 0.6745 |
| | WT vs. MERS-CoV | 0.5613 |
| AUC post-boost RBD IgG titers ($log_{10}$) Soluble 3M-052/Alum | WT vs. BA.4/BA.5 | 0.0034 |
| | WT vs. SARS-CoV-1 | 0.5957 |
| | WT vs. MERS-CoV | 0.5063 |
| AUC post-boost RBD IgG titers ($log_{10}$) RBD-LNH | WT vs. BA.4/BA.5 | 0.7716 |
| | WT vs. SARS-CoV-1 | 0.6382 |
| | WT vs. MERS-CoV | 0.9906 |

**Table S12.** *p* values for total GCBC, antigen-specific GCBC and Tfh counts elicited by a tetravalent betacoronavirus prime-boost of different vaccine formulations (general linear model followed by Student's t-test).

| | | Adjusted *p* values | |
|---|---|---|---|
| Total GCBCs ($log_{10}$) Week 2 | AddaVax vs. RBD-LNH | 0.0326 | * |
| | 3M-052/Alum vs. RBD-LNH | 0.3082 | ns |
| WT-specific GCBCs ($log_{10}$) Week 2 | AddaVax vs. RBD-LNH | 0.0044 | ** |
| | 3M-052/Alum vs. RBD-LNH | 0.0146 | * |
| Total Tfh cells ($log_{10}$) Week 2 | AddaVax vs. RBD-LNH | 0.0085 | ** |
| | 3M-052/Alum vs. RBD-LNH | 0.0178 | * |

**Table S13.** *p* values for percent of WT and BA.4/BA.5 antigen secreting cells (ASCs) isolated from bone marrow, elicited by a tetravalent betacoronavirus prime-boost of different vaccine formulations (general linear model followed by Student's t-test).

| | | Adjusted *p* values | |
|---|---|---|---|
| WT+ ASCs Week 2 | AddaVax vs. RBD-LNH | 0.2052 | ns |
| | 3M-052/Alum vs. RBD-LNH | 0.0442 | * |
| $BA.4/BA.5^+$ ASCs Week 2 | AddaVax vs. RBD-LNH | 0.6987 | ns |
| | 3M-052/Alum vs. RBD-LNH | 0.1249 | ns |
| $SARS\text{-}CoV\text{-}1^+$ ASCs Week 2 | AddaVax vs. RBD-LNH | 0.4864 | ns |
| | 3M-052/Alum vs. RBD-LNH | 0.0356 | * |
| $MERS\text{-}CoV^+$ ASCs Week 2 | AddaVax vs. RBD-LNH | 0.7563 | ns |
| | 3M-052/Alum vs. RBD-LNH | 0.1586 | ns |

**Table S14.** *p* values for anti-RBD endpoint IgG titers elicited in mice with pre-existing immunity vaccinated with a tetravalent betacoronavirus boost of different vaccine formulations (general linear model followed by Student's t-test).

| | | Adjusted *p* values | |
|---|---|---|---|
| Anti-WT RBD IgG titers ($log_{10}$) Week 2 post-boost | AddaVax vs. RBD-LNH | 0.0198 | * |
| | 3M-052/Alum vs. RBD-LNH | 0.0822 | ns |
| Anti-WT RBD IgG titers ($log_{10}$) Week 10 post-boost | AddaVax vs. RBD-LNH | 0.0518 | ns |
| | 3M-052/Alum vs. RBD-LNH | 0.1292 | ns |
| Anti-WT RBD IgG titers ($log_{10}$) Week 24 post-boost (M12) | AddaVax vs. RBD-LNH | 0.0339 | * |
| | 3M-052/Alum vs. RBD-LNH | 0.0142 | * |

| | | Adjusted *p* values | |
|---|---|---|---|
| Anti-BA.4/BA.5 RBD IgG titers ($log_{10}$) Week 2 post-boost | AddaVax vs. RBD-LNH | 0.0174 | * |
| | 3M-052/Alum vs. RBD-LNH | 0.0527 | ns |
| Anti-BA.4/BA.5 RBD IgG titers ($log_{10}$) Week 10 post-boost | AddaVax vs. RBD-LNH | 0.0052 | ** |
| | 3M-052/Alum vs. RBD-LNH | 0.0104 | * |
| Anti-BA.4/BA.5 RBD IgG titers ($log_{10}$) Week 24 post-boost (M12) | AddaVax vs. RBD-LNH | 0.0051 | ** |
| | 3M-052/Alum vs. RBD-LNH | 0.0007 | *** |

| | | Adjusted *p* values | |
|---|---|---|---|
| Anti-SARS-CoV-1 RBD IgG titers ($log_{10}$) Week 2 post-boost | AddaVax vs. RBD-LNH | 0.0227 | * |
| | 3M-052/Alum vs. RBD-LNH | 0.2056 | ns |
| Anti-SARS-CoV-1 RBD IgG titers ($log_{10}$) Week 10 post-boost | AddaVax vs. RBD-LNH | 0.0281 | * |
| | 3M-052/Alum vs. RBD-LNH | 0.8132 | ns |
| Anti-SARS-CoV-1 RBD IgG titers ($log_{10}$) Week 24 post-boost (M12) | AddaVax vs. RBD-LNH | 0.0448 | * |
| | 3M-052/Alum vs. RBD-LNH | 0.0653 | ns |

| | | Adjusted *p* values | |
|---|---|---|---|
| Anti-MERS-CoV RBD IgG titers ($log_{10}$) Week 2 post-boost | AddaVax vs. RBD-LNH | 0.7759 | ns |
| | 3M-052/Alum vs. RBD-LNH | 0.0655 | ns |
| Anti-MERS-CoV RBD IgG titers ($log_{10}$) Week 10 post-boost | AddaVax vs. RBD-LNH | 0.1186 | ns |
| | 3M-052/Alum vs. RBD-LNH | 0.2683 | ns |
| Anti-MERS-CoV RBD IgG titers ($log_{10}$) Week 24 post-boost (M12) | AddaVax vs. RBD-LNH | 0.3052 | ns |
| | 3M-052/Alum vs. RBD-LNH | 0.7345 | ns |

**Table S15.** *p* values for the area under the curve (AUC) over six months post-boost (M12) of anti-RBD endpoint IgG titers elicited in mice with pre-existing immunity vaccinated with a tetravalent betacoronavirus boost of different vaccine formulations (general linear model followed by Student's t-test).

| | | Adjusted *p* values | |
|---|---|---|---|
| Anti-WT RBD IgG titers ($log_{10}$) AUC | AddaVax vs. RBD-LNH | 0.0002 | *** |
| | 3M-052/Alum vs. RBD-LNH | 0.0827 | ns |

| | | Adjusted *p* values | |
|---|---|---|---|
| Anti-BA.4/BA.5 RBD IgG titers ($log_{10}$) AUC | AddaVax vs. RBD-LNH | <0.0001 | *** |
| | 3M-052/Alum vs. RBD-LNH | 0.0019 | ** |

| | | Adjusted *p* values | |
|---|---|---|---|
| Anti-SARS-CoV-1 RBD IgG titers ($log_{10}$) AUC | AddaVax vs. RBD-LNH | 0.0003 | *** |
| | 3M-052/Alum vs. RBD-LNH | 0.2248 | ns |

| | | Adjusted *p* values | |
|---|---|---|---|
| Anti-MERS-CoV RBD IgG titers ($log_{10}$) AUC | AddaVax vs. RBD-LNH | 0.0009 | *** |
| | 3M-052/Alum vs. RBD-LNH | 0.4698 | ns |

**Table S16.** *p* values for the area under the curve (AUC) over six months post-boost (M12) comparing anti-RBD endpoint IgG titers in mice with pre-existing immunity vaccinated with a tetravalent betacoronavirus boost of different vaccine formulations across antigens (general linear model followed by Tukey's HSD multiple comparisons).

| | | Adjusted *p* values |
|---|---|---|
| AUC post-boost RBD IgG titers ($log_{10}$) Soluble AddaVax | WT vs. BA.4/BA.5 | 0.0565 |
| | WT vs. SARS-CoV-1 | 0.2585 |
| | WT vs. MERS-CoV | 0.0343 |
| AUC post-boost RBD IgG titers ($log_{10}$) Soluble 3M-052/Alum | WT vs. BA.4/BA.5 | <0.0001 |
| | WT vs. SARS-CoV-1 | 0.1068 |
| | WT vs. MERS-CoV | <0.0001 |
| AUC post-boost RBD IgG titers ($log_{10}$) RBD-LNH | WT vs. BA.4/BA.5 | 0.4844 |
| | WT vs. SARS-CoV-1 | 0.8578 |
| | WT vs. MERS-CoV | 0.0007 |

**Table S17.** *p* values for the half-life of antibody titers against WT, BA.4/BA.5 SARS-CoV-2 and SARS-CoV-1 strains elicited in mice with pre-existing immunity vaccinated with a tetravalent betacoronavirus boost of different vaccine formulations (general linear model followed by Student's t-test).

| | | Adjusted *p* values | |
|---|---|---|---|
| Anti-WT RBD IgG titers ($log_{10}$) | AddaVax vs. RBD-LNH | 0.0146 | * |
| | 3M-052/Alum vs. RBD-LNH | 0.0125 | * |

| | | Adjusted *p* values | |
|---|---|---|---|
| Anti-BA.4/BA.5 RBD IgG titers ($log_{10}$) | AddaVax vs. RBD-LNH | 0.2652 | ns |
| | 3M-052/Alum vs. RBD-LNH | 0.2426 | ns |

| | | Adjusted *p* values | |
|---|---|---|---|
| Anti-SARS-CoV-1 RBD IgG titers ($log_{10}$) | AddaVax vs. RBD-LNH | 0.6472 | ns |
| | 3M-052/Alum vs. RBD-LNH | 0.5258 | ns |

**Table S18.** *p* values for $NT_{50}$ neutralization titers elicited in mice with pre-existing immunity vaccinated with a tetravalent betacoronavirus boost of different vaccine formulations (general linear model followed by Student's t-test).

| | | Adjusted *p* values | |
|---|---|---|---|
| Neutralization $NT_{50}$ titers ($log_{10}$) WT - Week 41 | AddaVax vs. RBD-LNH | 0.1711 | ns |
| | 3M-052/Alum vs. RBD-LNH | 0.1345 | ns |
| Neutralization $NT_{50}$ titers ($log_{10}$) Omicron BA.4/BA.5/BA.5.2 - Week 41 | AddaVax vs. RBD-LNH | 0.1393 | ns |
| | 3M-052/Alum vs. RBD-LNH | 0.5389 | ns |
